# Essential role of proline synthesis and the one-carbon metabolism pathways for systemic virulence of *Streptococcus pneumoniae*

**DOI:** 10.1101/2023.08.03.550501

**Authors:** Elisa Ramos-Sevillano, Giuseppe Ercoli, José Afonso Guerra-Assunção, Modupeh Betts, Frederick Partridge, Vitor E. Fernandes, Emma Wall, Stephen B. Gordon, Daniela M. Ferreira, Rob Heyderman, Jeremy S. Brown

**Affiliations:** Centre for Inflammation and Tissue Repair, UCL Respiratory, Division of Medicine, University College London, Rayne Institute, London WC1E 6JF, UK; Great Ormond Street Institute of Child Health, University College London (UCL), London WC1N 1EH, UK; Research Department of Infection, Division of Infection and Immunity, University College London, Rayne Institute, London WC1E 6JF, UK; Faculdade de Medicina e Ciências Biomédicas and ABC-RI. Faro, Portugal; Malawi-Liverpool-Wellcome Trust Clinical Research Programme Blantyre, Malawi; Liverpool School of Tropical Medicine Liverpool, United Kingdom

**Keywords:** *Streptococcus pneumoniae*, proline synthesis, formate-tetrahydrofoalte ligase, stringent response, virulence

## Abstract

Previous virulence screens have indicated potential roles during *Streptococcus pneumoniae* infection for the one-carbon metabolism pathway component Fhs and proline synthesis mediated by ProABC. To define how these metabolic pathways affect *S. pneumoniae* virulence we have investigated phenotypes and transcription profiles of Δ*fhs* and Δ*proABC* strain mutants. *S. pneumoniae* capsular serotype 6B BHN418 Δ*fhs* and Δ*proABC* mutant strains were markedly reduced virulence in mouse models of systemic infection and pneumonia, but were still able to colonise the nasopharynx. Although the Δ*fhs* and Δ*proABC* mutant strains grew normally in complete media, both mutant strains had markedly impaired growth in chemically defined medium, human serum and human CSF. The Δ*proABC* strain also had impaired growth under conditions of osmotic and oxidative stress. When transferred to the serotype 2 D39 *S. pneumoniae* strain background, the Δ*fhs* mutation replicated the virulence and growth in serum phenotype of the BHN418 mutation. In contrast, the D39 Δ*proABC* mutant could cause septicaemia and grow in human serum, indicating the role of this genetic locus during virulence is strain-specific. In human sera the Δ*fhs* and Δ*proABC* mutants both had major derangements in global gene transcription affecting multiple but different metabolic pathways, identifying the corresponding *S. pneumoniae* metabolic functions affected by these genes under infection-related conditions. Our data demonstrate an essential role for the *S. pneumoniae* one-carbon metabolism and a strain-conditional role for proline biosynthesis for growth in physiological fluids and therefore systemic infection, and further demonstrate the vital importance of bacterial metabolism for disease pathogenesis.

**Importance:** Rapid adaptation to grow within the physiological conditions found in the host environment is an essential but poorly understood virulence requirement for systemic pathogens such as *Streptococcus pneumoniae*. We have now demonstrated an essential role for the one-carbon metabolism pathway and a conditional role depending on strain background for proline biosynthesis for *S. pneumoniae* growth in serum or CSF and therefore for systemic virulence. RNAseq data demonstrated that loss of one carbon metabolism or proline biosynthesis both have profound but differing effects on *S. pneumoniae* metabolism in human serum, identifying the metabolic processes dependent on each pathway during systemic infection. These data provide a more detailed understanding of the adaptations required by systemic bacterial pathogens in order to cause infection, and demonstrate that the requirement for some of these adaptations vary between strains from the same species and could therefore underpin strain variations in virulence potential.

## INTRODUCTION

*Streptococcus pneumoniae* is a common upper respiratory tract commensal found in 10-15% of adults and 40-95% of infants, but is also a frequent cause of pneumonia, septicaemia or meningitis and is responsible for approaching a million deaths a year of children under the age of 5 worldwide (1–3). *S. pneumoniae* has a wide variety of virulence factors that contribute to disease pathogenesis (4), including the polysaccharide capsule required for immune evasion (5) and multiple surface proteins also involved in immune evasion as well as adhesion to host cells (6–9). Another essential requirement for bacterial virulence is survival and replication under the physiological conditions found within the host. Rapid replication in the lung is essential for *S. pneumoniae* to establish pneumonia (10), and growth in serum differentiates *S. pneumoniae* from the closely related but less virulent bacterial species *Streptococcus mitis* (11). The physiological conditions in the mammalian host include growth at 37°C, a pH of 7.4, serum osmolality of around 285 mmol / kg, and a restricted availability of specific nutrients needed for bacterial replication including several cations and micronutrients (12, 13). As a consequence the virulence of *S. pneumoniae* is dependent on cations, polyamine, and amino acid transporters (14–19), effective osmoregulation (18, 20), and synthesis of specific nutrients with limited availability in the host (21–23). However, at present our understanding of the *S. pneumoniae* factors required to replicate under physiological conditions remains very incomplete.

We analysed previously published transcriptome and transposon mutational screen analyses to identify metabolic pathways that are highly transcribed in mouse models of infection but whose role during infection has yet to be characterised in detail (24–26). From these studies we identified two genetic loci likely be involved in metabolic adaptation to growth in the host and mutation of which reduced virulence in mouse models (24), the *proABC* (SP0931-33) operon and *fhs* (SP1229). *proABC* is predicted to encode the enzymes ProA, a γ-glutamyl phosphate reductase, ProB, a γ-glutamyl kinase, and ProC, a pyrroline-5-carboxylate reductase responsible for proline synthesis from glutamate in a range of microorganisms (27). Proline protects bacteria against osmostress (28–30), and proline synthesis and / or transport is important for the virulence of several bacterial pathogens, including *Salmonella typhimurium* and *Mycobacterium tuberculosis* (31, 32). Virulence screens have shown that mutations affecting *proABC* reduces *S. pneumoniae* virulence in mouse models of pneumonia and meningitis (24, 33, 34).

*fhs* is predicted to encode a formate-tetrahydrofolate ligase that is part of the one-carbon metabolism pathway and catalyses the formation of 10-formyl-tetrahydrofolate from folate (as tetrahydrofolate, THF) and formate. The one-carbon metabolism pathway provides cofactors for the synthesis of multiple products, with THF donating a single carbon to the synthesis of amino acids and purines (35, 36). THF may also contribute to synthesis of the alarmones guanosine-pentaphosphate and - tetraphosphate [(p)ppGpp] that initiate the bacterial stringent response to adapt to nutritional and other physiological stresses (37). In many bacteria the synthesis of THF is usually performed by the FolD enzyme, but a minority of bacteria including *S. pneumoniae* use Fhs as an alternative mechanism (38–41). The one-carbon metabolism pathway could be important for multiple metabolic pathways involved in adaptation to host physiological conditions. Compatible with this, *fhs* expression was increased when *S. pneumoniae* were cultured in defined media containing low levels of methionine and during infection in a mouse model of meningitis (26, 35), and Rosconi et al. demonstrated that *fhs* is an essential gene required for growth in laboratory medium for a proportion of *S. pneumoniae* strains (37). Furthermore, mutation of *fhs* reduced *S. pneumoniae* virulence in mouse models of pneumonia or meningitis (24) (26), but the exact role of *fhs* duing infection has not been investigated. Defining the role of *fhs* during *S. pneumoniae* virulence could be relevant for multiple other bacterial pathogens whose genomes contain *fhs* (eg *Streptococcus pyogenes*, *Streptococcus agalactiae*, *Staphylococcus aureus*, and *Neisseria meningitidis)* (41).

Previously we have used *S. pneumoniae* BHN418 capsular serotype 6B mutant strains containing deletions of *proABC* or *fhs* in combination with deletion of the *pia* iron transporter locus as live attenuated *S. pneumoniae* vaccines, demonstrating the potential clinical utility of these genetic loci (42, 43). In this study we used *in vitro* and *in vivo* characterization of the *S. pneumoniae* Δ*proABC* and Δ*fhs* mutant strains to determine the roles of proline synthesis and the one-carbon metabolism pathway during disease pathogenesis.

## RESULTS

### Bioinformatic analysis and mutation of *fhs* and *proABC*

The predicted protein product for the *S. pneumoniae* TIGR4 strain *proA* (Sp_0932), *proB* (Sp_0931), and *proC* (Sp_0933) has 48% 42% and 28% amino acid sequence identity respectively to the ProA, ProB, and ProC proteins from *Bacilus subtilis* (strain 168), and 46%, 38% and 40% from *Escherichia coli* (strain K12) (44). The predicted amino acid sequences for ProA, ProB and ProC have 96%, 98% and 75% identity respectively compared to the TIGR4 sequence across 18 *S. pneumoniae* strains. The predicted protein product for the *S. pneumoniae* TIGR4 strain *fhs* (Sp_1229) Fhs has 43% amino acid sequence identity to *Escherichia coli* Fhs (44), and is 99-100% conserved at the amino acid level when 13 *S. pneumoniae* strains were compared to the predicted TIGR4 sequence. All the described active sites of Fhs were found in the predicted sequence of *S. pneumoniae* Fhs, including an ATP-binding domain (PTPAGEGKXT, where X is either S or T), a glycine (G)-rich nucleotide binding consensus sequence, and residues involved in binding to folate (Trp412 or Phe 385), para-aminobenzoic acid (PABA) (Pro385 or Leu408), or THF (residues 95 to 103 EPSLGPX_2_G with an aspartate at residue 29) (36) (45, 46) (47). Bioinformatics analysis using PSIPRED (48) indicated that Fhs is located intracellularly. For phenotpye analyses, mutant strains containing complete deletion of *proABC* and *fhs* were constructed in the capsular serotype 6B strain BHN48 using overlap extension PCR (**Fig. 1A and B**) and transferred to the capsular serotype 2 D39 strain using transformation with genomic DNA. A Δ*fhs*+*fhs* serotype 6B complemented mutant was constructed by insertion of a native copy of *fhs* between the 423898 to 423940 and 423358 to 423394 sites (D39 strain base pair numbers) in the genome using the promoterless integration vector pPEPY (49).

**Figure 1.**
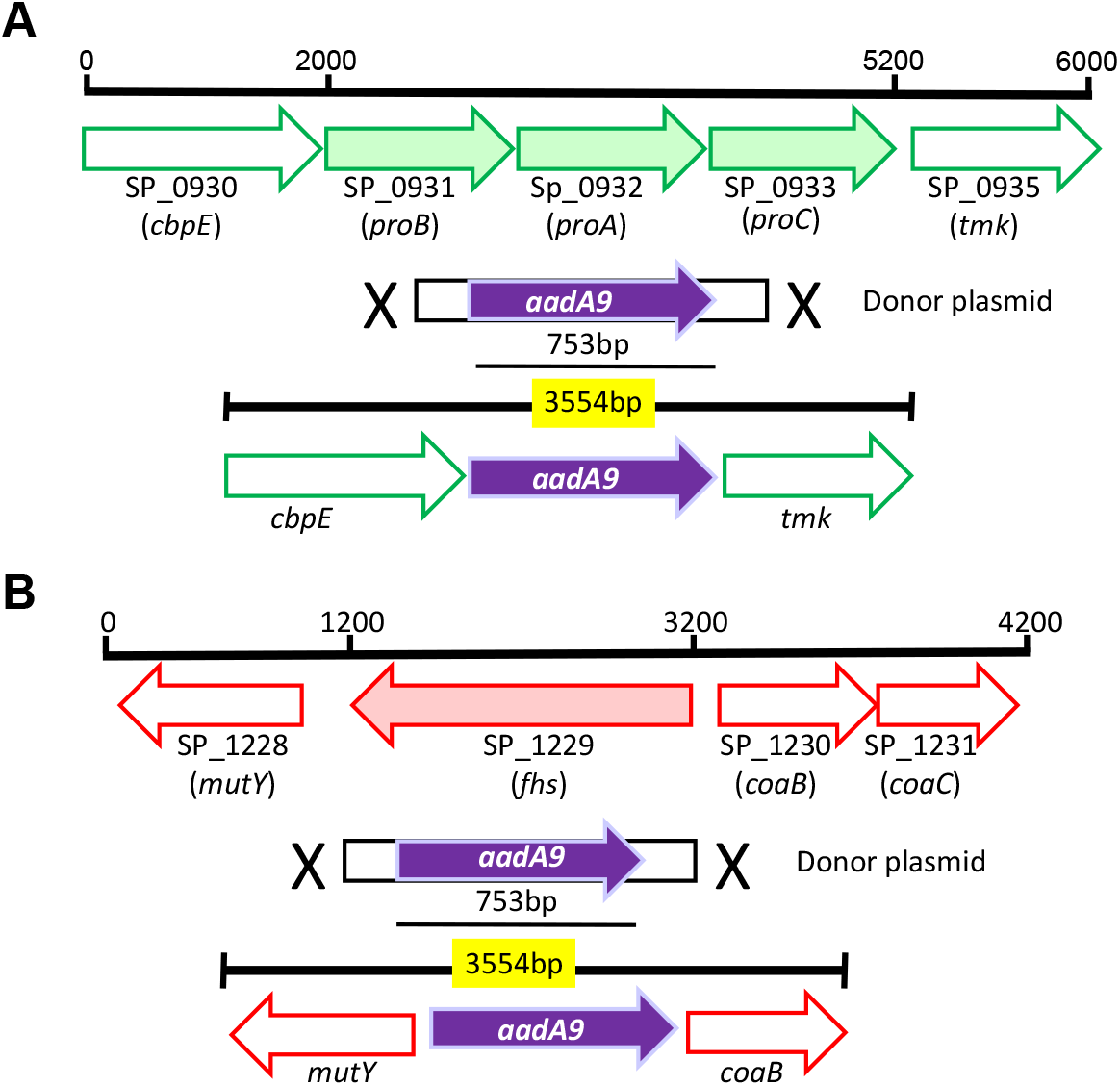
Genetic organization and construction of *proABC* and *fhs* deficient strains in 6B BHN418 strain of *S. pneumoniae*. (**A**) Schematic of the Sp_0931-0933 locus with the TIGR4 genome gene number and the assigned gene names in parentheses, if available. Arrows indicate transcriptional direction. The putative *proABC* operon is shaded in green colour. Structure of the Sp_0931-33 locus in the Δ*proABC* mutant strain is replaced with an in-frame copy of spectiniomycin (*aadA9*), which is shaded purple. (**B**) Schematic of the Sp_1229 locus with the TIGR4 genome gene number and the assigned gene names in parentheses, if available. Arrows indicate transcriptional direction. The putative *fhs* gene is shaded in red colour. Structure of the Sp_1229 gene in the Δ*fhs* mutant strain is replaced with an in-frame copy of spectiniomycin (aadA9), which is shaded purple.

### Δ*proAB*C and Δ*fhs* strain *in vivo* phenotypes

The virulence of the 6B *Δfhs* or *ΔproABC* strains were compared to the wild-type 6B strain in mouse models of pneumonia, septicaemia, and colonisation. As the capsule is the most important *S. pneumoniae* virulence factor, the unencapsulated Δ*cps* 6B strain was used to assess the relative strength of any virulence defect of the *Δfhs* or *ΔproABC* strains. In the mouse model of pneumonia both the Δ*proABC* and Δ*fhs* 6B mutant strains had a similar phenotype to the Δ*cps* unencapsulated strain, with almost complete failure to disseminate from the lungs to the blood (**Fig. 2A**) and non-significant reductions in lung CFU compared to the wild-type strain (**Fig. 2B**). In the sepsis model, the Δ*proABC* and Δ*fhs* mutant strains showed large reductions in blood or spleen CFU compared to the wild-type strain 6B (**Fig. 3A and 3B**). In both the pneumonia and sepsis infection models, target organ CFU were similar for mice infected with the complemented 6B Δ*fhs*+*fhs* mutant strain to those infected with the wild-type (**Fig. 2C and D, 3C and D**), confirming the virulence defect observed for Δ*fhs* strain was due to deletion of *fhs*. When transferred into the D39 *S. pneumoniae* strain background, a similar virulence phenotype to the 6B strain was observed for the D39 Δ*fhs* strain in both pneumonia (**Fig. 2E and 2F**) and sepsis models (**Fig. 3E and 3F**) and the D39 Δ*proABC* strain in the pneumonia model (**Fig. 2E and 2F**). However, in the sepsis model the Δ*proABC* D39 strain virulence was relatively preserved compared to the Δ*proABC* 6B strain, with a statistically non-significant reduction of approximately 2 log_10_ in blood and spleen CFU (**Fig. 3E and 3F**). In contrast to the sepsis and pneumonia models, in the nasopharyngeal colonisation model nasal wash CFU for the 6B Δ*proABC* and Δ*fhs* strains were maintained at similar levels to the wild-type strain at 7 days and approximately 2 log_10_ greater than nasal wash CFU for mice infected with the Δ*cps* strain (**Fig. 2G**). The Δ*proABC* and Δ*fhs* 6B strains maintained colonisation of the nasopharynx up to 12 days, although with significant reductions in nasal wash CFU compared to wild-type at this later timepoint (**Fig. 2H**). To confirm the differences in target organ CFU were associated with reductions in lethality, the development of disease was monitored for 7 days in the pneumonia model after infection with the 6B Δ*proABC* or Δ*fsh* mutant strains. 50% of the mice inoculated with wild-type 6B strain or the complemented Δ*fhs* mutant developed fatal infection over a period of 72 hours (**Fig. 3G**), whilst 90% of mice infected with Δ*proABC* and all mice infected with *Δfhs* mutant strains survived. These data demonstrate that loss of *fhs* has a profound effect on systemic virulence in both 6B and D39 backgrounds, whereas the effects of loss of *proABC* on virulence was strain dependent. Despite loss of systemic virulence, both mutant strains were able to colonise the nasopharynx for a prolonged period.

**Figure 2.**
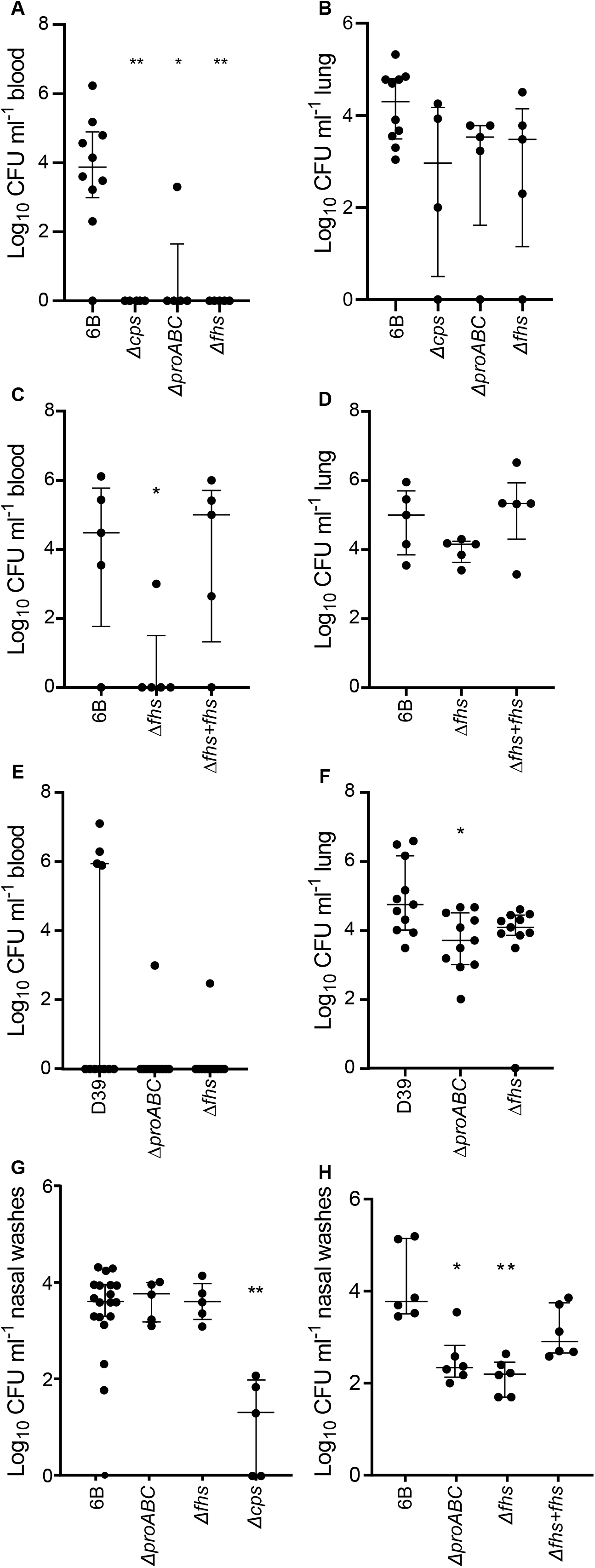
Assessment of virulence of the Δ*fhs* and Δ*proABC* mutants strains on a pneumonia and colonsation models. Log_10_ ml^-1^ bacteria CFU recovered from blood (**A**, **C**, **E**) and lung (**B**, **D**, **E**) of five week old CD-1 mice infected via intranasal route after 18 hours post inoculation with 1×10^7^ CFU of the wild-type 6B or D39 and mutant strains Δ*proABC* and Δ*fhs*. Each symbol represents CFU data from a single mouse, horizontal bars represent median values, error bars represent interquartile range and asterisks represent statistical significance compared to the wild-type strain (Kruskal–Wallis with Dunn’s post hoc test to identify significant differences between groups, * *p* < 0.05; ** *p* < 0.01). (**G**,**H**) Colonisation model; CFU in nasal washes of CD1 mice 7 days(G) or 12 days (**H**) post colonisation with 1×10^7^_CFU of wild-type 6B or single mutant *S. pneumoniae* strains.

**Figure 3.**
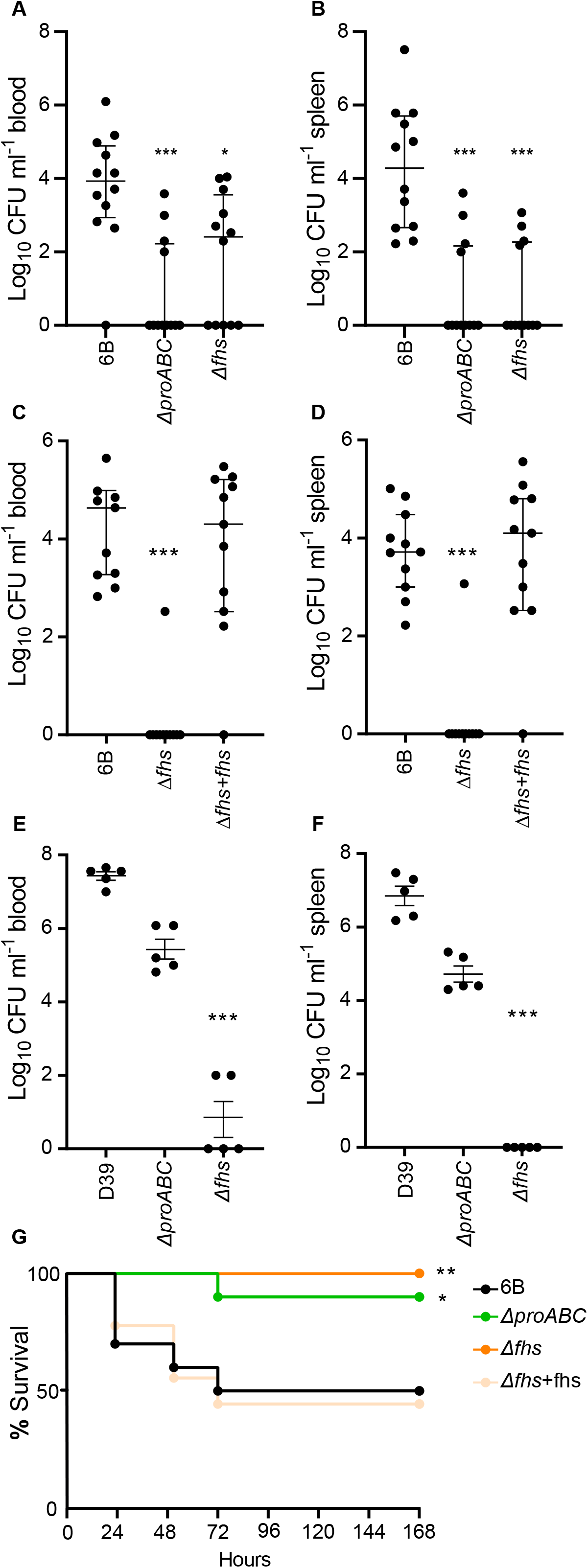
Assessment of virulence of the Δ*fhs* and Δ*proABC* mutants strains on a sepsis model and survival analysis of CD-1 mice during pneumococcal pneumonia. Log_10_ ml^-1^ bacteria CFU recovered from blood (**A, C, E**) and spleen (**B, D, F**) of five week old CD-1 mice infected via intraperitoneal route after 24 hours post inoculation with 5×10^6^ CFU of the wild-type (6B or D39) or mutant strains Δ*proABC*, Δ*fhs* and fhs complemented mutant strain Δ*fhs*+*fhs*. Each symbol represents CFU data from a single mouse, horizontal bars represent median values, error bars represent interquartile range and asterisks represent statistical significance compared to the wild-type strain (Kruskal–Wallis with Dunn’s post hoc test to identify significant differences between groups, * *p* < 0.05; ** *p* < 0.01; *** *p* < 0.001). (**G**) Survival of five week old CD-1 mice (n=10) infected via intranasal was monitored over a 7-day period. Survival curves were determined to be significantly different by Logrank (Mantel-Cox) test (* *p* < 0.05; ** *p* < 0.01).

### *S. pneumoniae fhs* and *proABC* were not required for immune evasion

Confocal microscopy using an anti-capsule antibody provided no evidence that loss of virulence of the 6B Δ*proABC* and Δ*fhs* strains were due to effects of the mutations on cell morphology or capsule thicknness (**Fig. 4A**). After incubation in human sera neither strain showed increased recognition by complement or antibody, or increased sensitivity to neutrophil killing (**Fig. 4B to F**). Furthermore in a nematode worm model of infection that reflects host toxicity caused by *S. pneumoniae* (50, 51) the Δ*fhs* mutant strains killed *C. elegans* as rapidly as the wild-type strain within 6 hours of infection (**Fig. 4G to H**). The Δ*proABC* mutant did show some delay in killing with 100% of the worms killed only after 24 hours (**Fig. 4H**) suggesting there was some effect of loss of *proABC* on host toxicity in this model. Overall these data indicate the reduced virulence of the Δ*proABC* and Δ*fhs* mutant strains was not related to increased susceptibility to host immunity.

**Figure 4.**
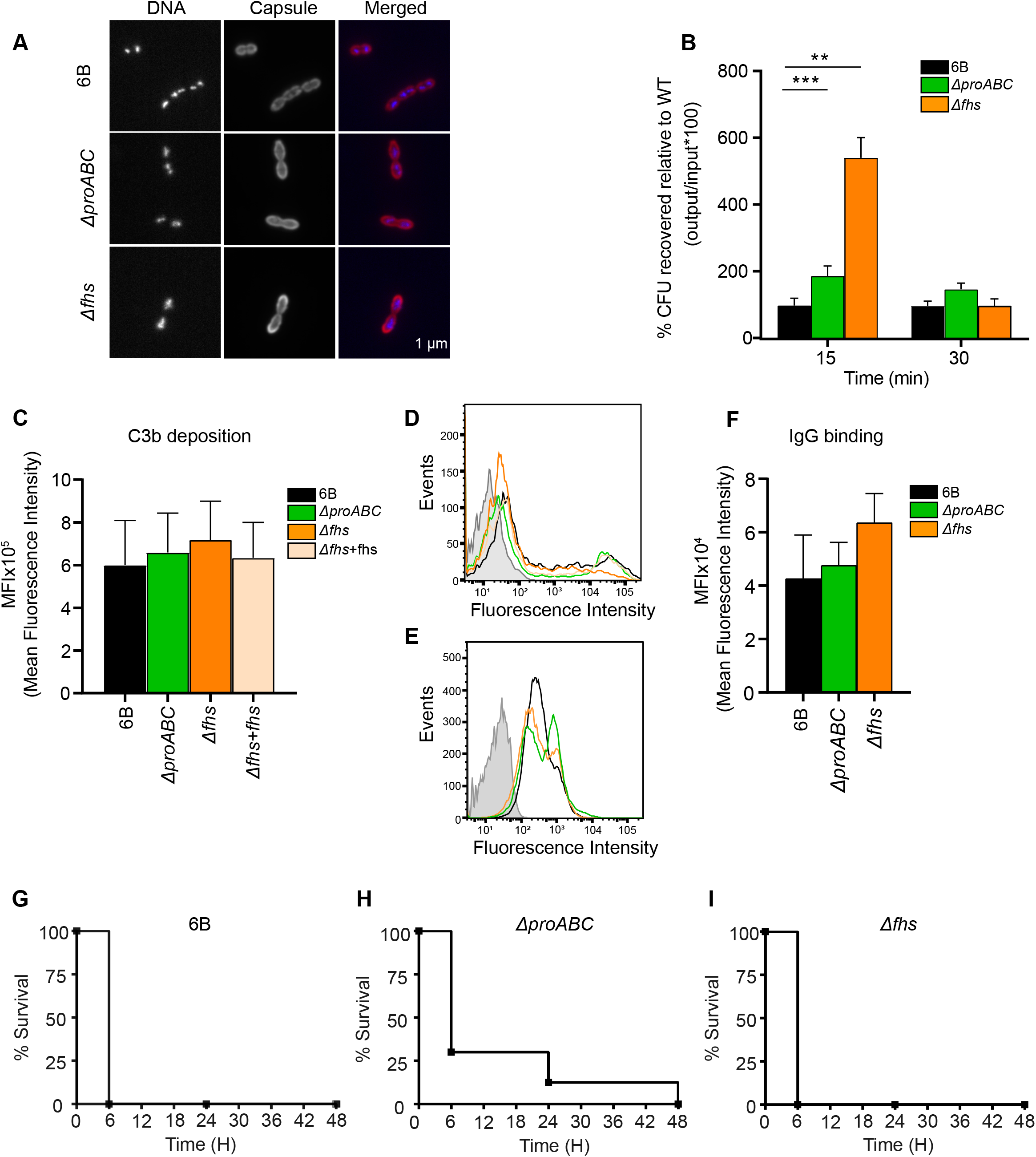
Effect of *fhs* and *proABC* absence in host immunity. (**A**) Fluorescent microscopy of wild-type and mutant strains following incubation with 4′,6-diamidino-2-phenylindole (DAPI) (binds to DNA to identify bacterial cells, panels in first column) or pneumococcal antiserum labelled with Alexa fluor 546 (recognizes serotype 6 capsule, panels in second column). The scale bar (bottom right) represents 1 µm. (**B**) Bacterial survival in a neutrophil killing assay (the multiplicity of infection or MOI is 1 bacterium:100 neutrophils) represented as %CFU recovered after 15 or 30 min incubation relative to the wild-type bacteria. (**C**) Mean fluorescence Intensity (MFI) of C3b deposition measured using a flow cytometry assay on the 6B strain and the mutant strains Δ*fhs*, complemented strain Δ*fhs*+fhs and Δ*proABC* in 25% of human serum. (**D**) Example of flow cytometry histogram for C3B deposition. Grey shading indicates the results for bacteria incubated in PBS alone. (**E**) Example of flow cytometry histogram for IgG binding. Grey shading indicates the results for bacteria incubated in PBS alone (**F**) MFI of IgG binding measured using a flow cytometry assay on the 6B strain and the mutant strains Δ*fhs* and Δ*proABC* in 25% of human serum. (**G**,**H**,**I**) *S. pneumoniae* kills *C. elegans* in a solid assay. Nematodes were fed on *S. pneumonaie* 6B strain or mutant strains Δ*proABC* and Δ*fhs*. Both (**G**) wild-type *S. pneumoniae* 6B and (**I**) Δ*fhs* mutant strains killed all nematodes within 6 h, whereas (**H**) Δ*proABC* mutant strain showed intermediate or poor killing. Similar results were obtained in 3 independent experiments.

### Growth of **Δ***proABC* and **Δ***fhs* strains under conditions of osmotic, cation and oxidative stress

Culture in laboratory media was used to investigate whether Δ*proABC* and Δ*fhs* strains had impaired growth under specific physiological stress conditions. In rich media (THY) the Δ*proABC* and Δ*fhs* 6B strains had identical growth to the wild-type strain (**Fig. 5A and 6A**). Addition of NaCl or sucrose to THY to induce osmotic stress or paraquat to cause oxidative stress resulted in impaired growth of the Δ*proABC* strain compared to the wild-type (**Fig. 5B to D**) (44, 52, 53), but did not consistently affect growth of the Δ*fhs* strain (**Fig. 6B** to **6D**). In cation-depleted THY, growth of both the Δ*proABC* and Δ*fhs* strains was slightly impaired compared to the wild-type strain (**Fig. 5E and 6E)**.

**Figure 5.**
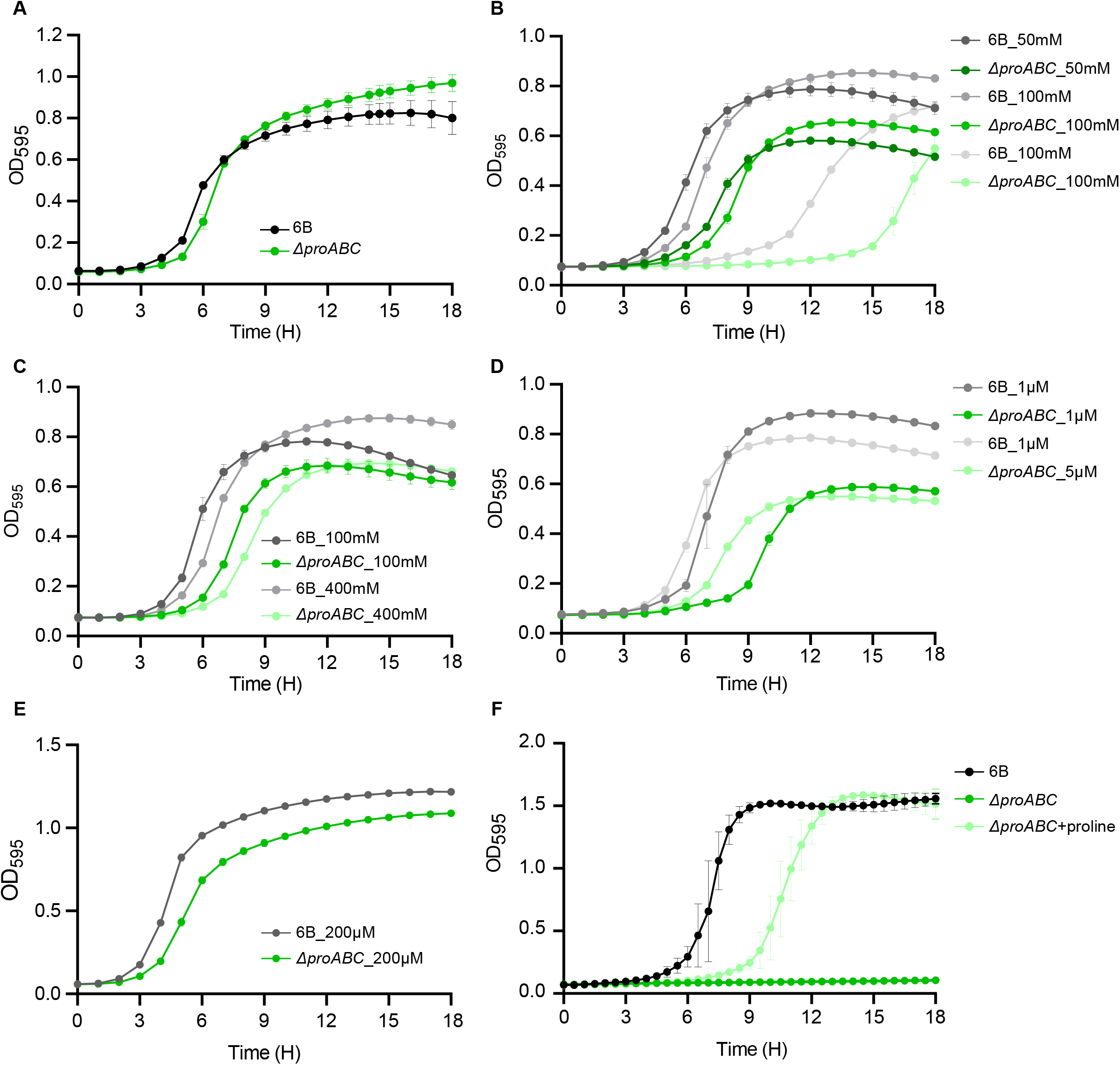
*In vitro* characterization of the Δ*proABC* mutant strain. (**A**) Growth of wild-type 6B and Δ*proABC* strains in THY broth. (**B**) Growth of wild-type 6B and Δ*proABC* strains in THY broth supplemented with 50, 100 and 200mM of NaCl. (**C**) Growth of wild-type 6B and Δ*proABC* strains in THY broth supplemented with 100 and 400mM of sucrose. (**D**) Growth of wild-type 6B and Δ*proABC* strains in THY broth supplemented with 1 and 5mM of paraquat. (**E**) Growth of wild-type 6B and Δ*proABC* strains in THY broth supplemented with 200uM of Ethylenediamine-N,N′-diacetic acid (EDDA). (**F**) Growth of wild-type 6B and Δ*proABC* strains in CDM media. Growth in all conditions was assessed at 37°C and 5% CO_2_ every 30 minutes for a period of 24 hours by using a plate reader and measuring OD_595_.

**Figure 6.**
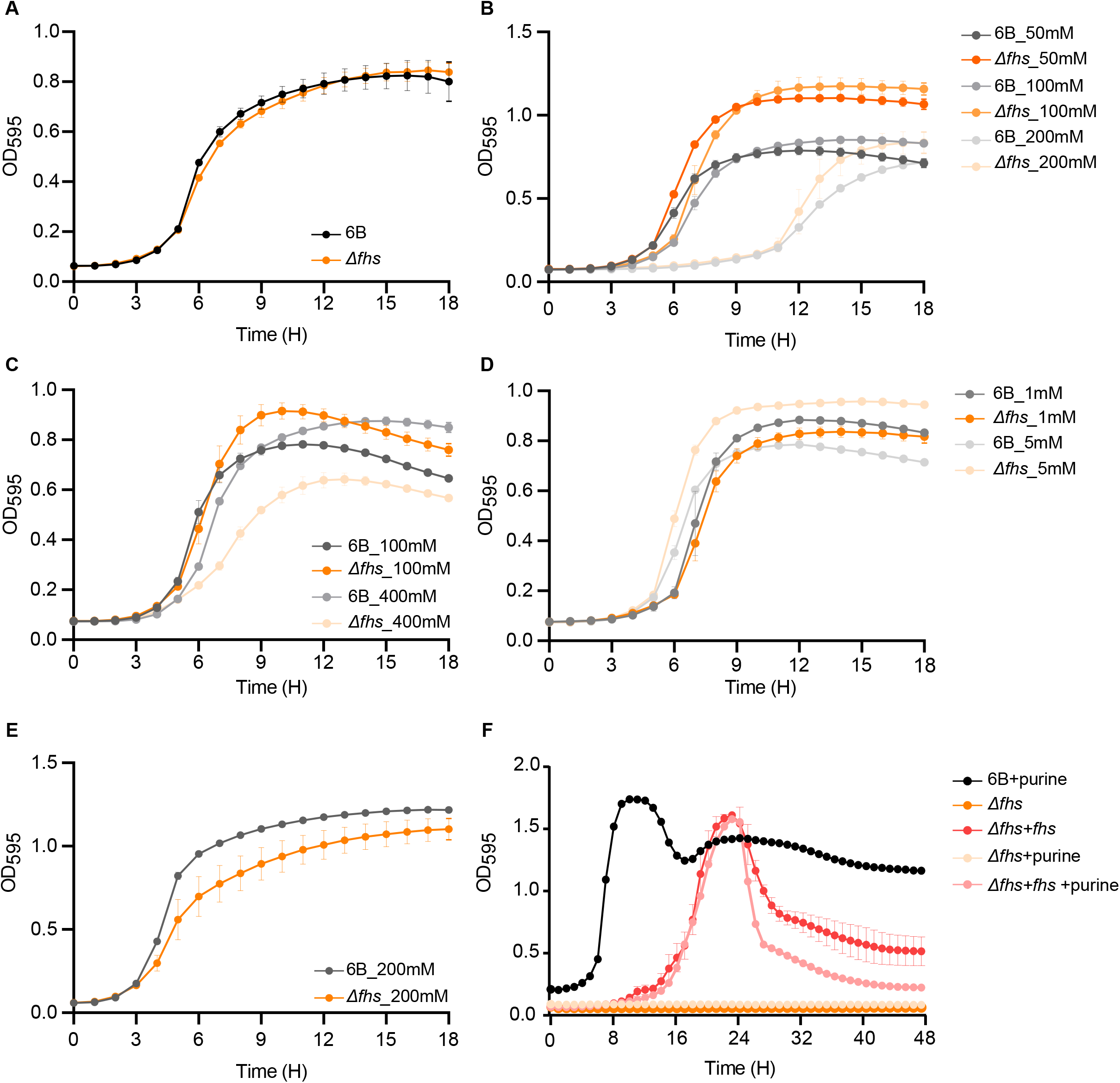
*In vitro* characterization of the Δ*fhs* mutant strain. (**A**) Growth of wild-type 6B and Δ*fhs* strains in THY broth. (**B**) Growth of wild-type 6B and Δ*fhs* strains in THY broth supplemented with 50, 100 and 200mM of NaCl. (**C**) Growth of wild-type 6B and Δ*fhs* strains in THY broth supplemented with 100 and 400mM of sucrose. (**D**) Growth of wild-type 6B and Δ*fhs* strains in THY broth supplemented with 1 and 5mM of paraquat. (**E**) Growth of wild-type 6B and Δ*fhs* strains in THY broth supplemented with 200uM of Ethylenediamine-N,N′-diacetic acid (EDDA). (**F**) Growth of wild-type 6B, Δ*fhs* and the complemented Δ*fhs*+*fhs* strains in CDM media supplemented with purine 1mg ml^-1^. Growth in all conditions was assessed at 37°C and 5% CO_2_ every 30 minutes for a period of 24 hours by using a plate reader and measuring OD_595_.

### Growth of the **Δ***proABC* and **Δ***fhs* strains is highly impaired in chemically defined media

Growth in the defined medium CDM using glucose as the carbon source was used to assess whether the 6B Δ*proABC* or Δ*fhs* mutations impaired bacterial replication under conditions with restricted availability of nutrients. Both the Δ*proABC* and Δ*fhs* strains had severe growth defects in CDM media compared to the wild-type parental strain (**Fig. 5F and 6F**). Growth of the Δ*proABC* strain in CDM was restored by supplementation with 1mg ml-1 proline alone (**Fig. 5F),** but not by addition of proline containing peptides (at 10, 50 or 250 ug/ml) imported by the *S. pneumoniae* AliA and AliB ABC transporters or by an oligopeptide consisting of 8 proline residues (54) (**Suppl Fig. 2**). These data demonstrate environmental proline can compensate for loss of proline synthesis by *S. pneumoniae* due to deletion of *proABC*, but this is probably dependent on proline-specific uptake transporters rather than oligopeptide transporters. Despite the likely role of Fhs in purine synthesis, addition of purine, adenine, formate or glycine as supplements for growth in minimal medium (previously suggested to compensate for loss of growth for a Δ*folD/p-fhs E. coli* strain (39)) did not restore growth of the *fhs* mutant strain in minimal medium CDM (**data not shown**, **Fig. 6F**). Overall, these results indicate that replication of the *S. pneumoniae* 6B strain in conditions with limited availability of multiple micro- and macronutrients is highly dependent on both proline synthesis and the *S. pneumoniae* one-carbon metabolism pathway.

### **Δ***fhs* and **Δ***proABC* mutant strains grew poorly in physiological fluids

The results of the above experiments suggested the reduced virulence of the *S. pneumoniae* 6B Δ*fhs* and Δ*proABC* strains could be related to poor growth in the physiological conditions found in the host. To investigate this, growth of both mutant strains was compared to the wild-type strain in *ex vivo* human sera or CSF. The 6B Δ*proABC* mutant was markedly attenuated in growth in human sera and completely attenuated in growth in human CSF (**Fig. 7A and 7B**). The supplementation of fresh sera or CSF with two different concentrations of proline (0.1 and 1 mg ml^-1^) substantially improved Δ*proABC* growth, linking the poor growth of this strain directly to proline availability (**Fig. 7A and 7B**). The Δ*fhs* mutant also had markedly impaired growth in both human sera and CSF, which was restored for the Δ*fhs+*fhs complemented strain (**Fig. 7C and D**). In contrast to the CDM data, the growth of the Δ*fhs* mutant in human serum was also partially complemented by addition of purine (**Fig. 7E**). In a laboratory media that mimicks fluid nasal (55), the Δ*proABC* had a similar growth profile to the wild-type strain whereas the Δ*fhs* mutant had markedly reduced growth (**Fig 7F**). To assess the potential effects of strain background on the *fhs* and *proABC* roles during infection, the growth of the D39 Δ*fhs* and Δ*proABC* mutant strains in serum was investigated. The D39 Δ*fhs* strain had delayed growth in serum, although this strain eventually reached a similar maximum OD to the wild-type strain (**Fig. 7G**). In contrast to the results for the 6B strain, the D39 Δ*proABC* mutation grew normally in serum, a result that is compatible with the ability of this strain to still cause septicaemia in the mouse sepsis model. Overall, these data link the markedly impaired systemic virulence of *S. pneumoniae* 6B and D39 Δ*fhs* and 6B Δ*proABC* strains to their poor replication in physiological fluids, with an essential role for disease pathogenesis for the one-carbon metabolism pathway whereas the role of proline synthesis was strain-dependent.

**Figure 7.**
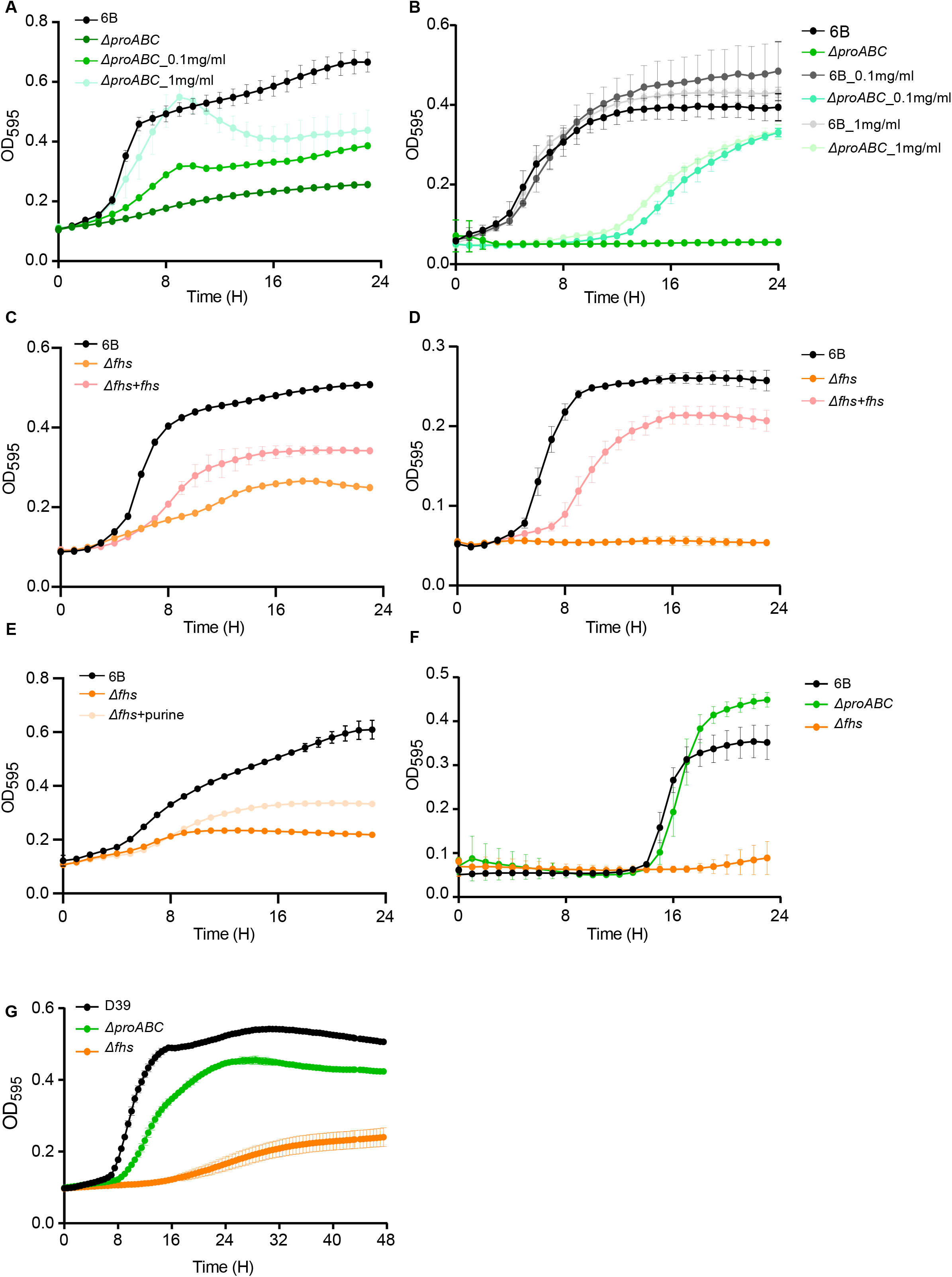
Phenotypic characterization of Δ*proABC* and Δ*fhs* mutant strains in biological fluids. (**A**) Growth of wild-type 6B and Δ*proABC* mutant strains in human serum supplemented with proline 0.1mg ml^-1^ and 1mg ml^-1^. (**B**) Growth of wild-type 6B and Δ*proABC* mutant strain in human cerebrospinal fluid (CSF) supplemented with proline 0.1mg ml^-1^ and 1mg ml^-1^. (**C**) Growth of wild-type 6B,Δ*fhs* mutant and complemented Δ*fhs+fhs* strain in human serum or (**D**) in CSF for a 24 hours period. (**E**) Growth of wild-type 6B and Δ*fhs* mutant strain in human serum supplemented with purine 1 mg ml^-1^. (**F**) Growth of wild-type 6B and mutant strains Δ*proABC* and Δ*fhs* in nose like media (The main carbon source of this media is N-Acetylglucosamine). (**G**) Growth of wild-type D39 and Δ*proABC* mutant strains in human serum. Growth in all conditions was assessed at 37°C and 5% CO_2_ every 30 minutes for a period of 24 hours by using a plate reader and measuring OD_595_.

### RNA-seq of *S. pneumoniae* 6B strains cultured in serum and THY

To characterise *S. pneumoniae* adaptations during growth under physiological conditions found during invasive infection and how these were affected by the Δ*proABC* and Δ*fhs* mutations, RNA-seq was performed on biological replicates of the 6B wild-type, Δ*proABC,* and Δ*fhs* 6B strains grown to mid-exponential phase in THY before incubation for a further 60 minutes in human sera or THY. Principal component analysis of normalised and transformed transcripts showed clear separations between all three strains in serum, with less clear separation when grown in THY (**Suppl. Fig.2**). The first two principal components (PC) accounted for about 90% of the variability between mutant and wild-type strains, with 66% of the total variance in normalized gene expression between the strains accounted for by the PC1. Selected operons showing changes in expression by the wild type 6B strain between incubation in serum or THY are shown in **Table 1** and **Suppl. Table 2**. Both the Δ*proABC* and Δ*fhs* mutant strains showed increased or decreased expression of a similar number of genes when incubated in THY compared to the wild type strain (**Fig. 8A and 8D**). Changes in the Δ*proABC* transcriptome compared to the wild-type strain in THY were dominated by genes involved in carbohydrate utilisation and biosynthesis, with increased expression of the maltose / maltodextrin ABC transporter and the mannose and glucose phosphotransferase (PTS) transporters (**Figure 8B, Suppl. Table 3**) (56). When cultured in THY the Δ*fhs* strain upregulated operons affecting multiple biochemical functions, including amino acid metabolism and synthesis, the Piu iron uptake ABC transporter, a PTS sugar transporter, and other aspects of metabolism (**Figure 8E, Suppl. Table 3**). In THY both the Δ*fhs* and Δ*proABC* strains upregulated genes involved in fatty acid synthesis and downregulation of the chaperon proteins *groEL, dnaJK,* and the chaperon regulator *hrcA* (**Fig. 8B and 8E**), perhaps due to non-specific responses to metabolic stress caused by the mutations. These data show that despite the maintenance of normal growth in THY by the Δ*proABC* and Δ*fhs* strains both mutations were still associated with significant changes in gene expression likely to due to bacterial adaptation to loss of specific biochemical functions related to each mutation.

**Figure 8.**
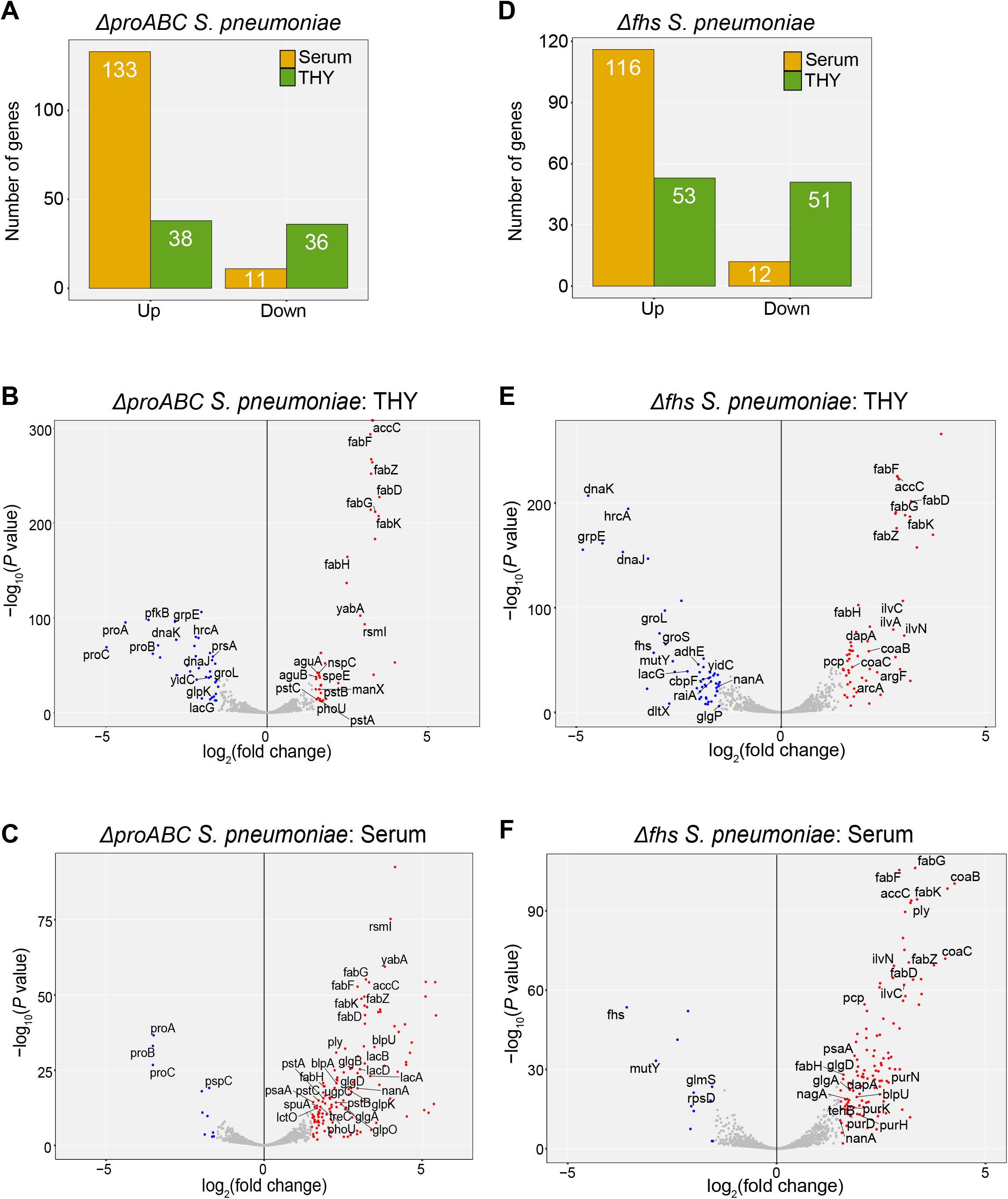
Transcriptomic changes of mutant strains in THY and human serum compared to wild-type *S. pneumoniae*. (**A**) Total number of genes for which expression was significantly up- or down-regulated genes by the Δ*proABC* strain compared to wild-type in THY or human serum. Differentially expressed genes (DEGs) were defined as those with log_2_ fold change of > 1.5 and FDR of < 0.05. Volcano plots showing the individual profiles of DEGs for the Δ*proABC* strain grown in (**B**) THY or (**C**) human serum (**D**) Total number of genes for which expression was significantly up- or down-regulated genes by the Δ*fhs* strain compared to wild-type in THY or human serum. Volcano plots showing the individual profiles of DEGs for the Δ*fhs* strain grown in (**E**) THY or (**F**) human serum. Red dots represent DEGs upregulated in the mutant strain, and blue dots represent downregulated DEGs. Grey coloured dots are transcripts that did not meet logFC and FDR thresholds for DEGs. Short gene names of DEGs are annotated on the plot when available. Data are from 3 biological replicates.

**Table 1:** Adjacent genes and operons showing differential expression (log_2_ fold change) by the wild-type 6B strain when cultured in sera compared to THY.

| Strain gene numbers and category |  |  | Gene names | Function | log<sub>2</sub> RNAseq ratio in sera<br>versus THY |
| --- | --- | --- | --- | --- | --- |
| 6B BHN418 | TIGR4 | D39 |  |  |  |
| Amino acid uptake and metabolism |  |  |  |  |  |
| Spn_00425-29 | SP2116-20 | SPD1945-49 |  | CAAX amino terminal protease family protein | -2.13,-2.56,-3.14,-3.39,-3.06 |
| Spn_00434-36 | SP2125-2126 | SPD1954-56 |  | branched-chain amino acid biosynthetic pathway | -2.79,-2.80,-4.01 |
| Spn_00659-60 | SP0112-13 | SPD0109-10 | *artP1, argG* | Arginino-succinate synthase | -3.11,-3.20 |
| Spn_00839-44 | SP0275-80 | SPD0255-60 | *?, yafQ, _polC_2, pepS, ? rsuA1* | Cleavage of amino acid | -3.53,-3.11,-2.77,-3.1,-2.53,-1.6 |
| Spn_01301-05 | Spn_0749-53 | SP0750-54 | *livJ, livH, livM, lptB, livF* | BCAA* ABC transporter | -3.38,-2.26,-2.03,-1.87,-2.19 |
| Spn_01699 | SP1159 | SPD1023 | *xerS* | tyrosine recombinase | -3.48 |
| **Sugar uptake and metabolism** |  |  |  |  |  |
| Spn_00150-52 | SP1882-84 | SPD1662-64 | *treC, treP, treR* | Sucrose metabolism | 4.55,4.50,2.20 |
| Spn_01423-25 | SP0875-77 | SPD0771-3 | *fruR, fruB, fruA* |  | 5.28,4.99,4.69 |
| **Other metabolism** |  |  |  |  |  |
| Spn_00128 | SP1859 | SPD_1640 | *pnuC* | nicotinamide mononucleotide transporter | -6.57 |
| Spn_0136-39 | SP1869-72 | SPD1649-52 | *feuB, fepD1, FHUc, yclQ* | Iron transport | 1.78,1.81,1.77,1.79 |
| Spn_00311 | SP2016 | SPD_1826 | *nadC* | nicotinate-nucleotide pyrophosphorylase | -4.44 |
| Spn00603-7,<br>00609, 00611 | SP0044-48,<br>50, 53 | SPD0051-55,<br>0057, 0059 | *purC, purI, purF, purM, purN, purH, purE* | Purine / biotin / coenzyme A synthesis | -2.31,-3.44,-2.68,-2.71,-2.32-2.26-1.74 |
| Spn_01501-02 | SP0963-64 | SPD0851-52 | *pyurK, pyrDb* |  | -4.46,-3.86 |
| Spn_01807-10 | SP1275-78 | SPD1131-34 | *carB, carA, pyrB, pyrR* | Pyrimidine synthesis | -3.50,-3.82,-3.63,-3.40 |
| Spn_00482-85 | SP2173-76 | SPD_2002-6 | *dltD, dltC, dltB, dltA* | Cell wall synthesis | 3.18,2.82,2.95,2.76 |
| **Other** |  |  |  |  |  |
| Spn_00601-2 | SP0042-43 | SPD0049-50 | *comA, comB* | competence factor transport proteinS | -1.99,-1.80 |
| Spn_00698 | SP0141 | SPD_0144 | *mutR* | Positive transcriptional regulator of mutA | -4.15 |
| Spn_00914 | SP0366 | SPD_0334 | *aliA* | oligopeptide ABC transporter, | -5.16 |
| Spn_01082 | SP0517 | SPD0460 | *dnaK3* | Molecular chaperone | -4.31 |
| Spn_01631_pulA_2 | Sp1118 | SPD1002 | *pulA2* | pullulanase | 3.04 |
| Spn_02064 | SP1161 | SPD1025 | *lpd* | dihydrolipoamide dehydrogenase | 3.11 |

### Marked disruption of gene expression by the **Δ***proABC* and **Δ***fhs* strains in serum reflected different metabolic responses caused by each mutation

Culture in serum instead of THY resulted in a marked increase in the total number of genes showing increased expression compared to the wild-type strain for both the Δ*proABC* (133 versus 36 genes) and Δ*fhs* (116 versus 51 genes) strains (**Fig. 8A and 8D**), demonstrating that under infection-related conditions both mutant strains underwent major compensatory changes in gene expression patterns. Compared to the wild-type strain the Δ*proABC* strain showed increased expression in serum of ten operons involved in sugar uptake and metabolism (eg Sp_0060-64 and Sp_0645-48 PTS sugar transporter operons), as well as several other ABC transporters and four operons containing genes of unknown function (**Fig. 8C**, **Table 2, Suppl. Table 4)**. In contrast, the genes upregulated in sera by the Δ*fhs* strain included 4 operons involved in amino acid uptake or biosynthesis, several ABC transporters, teichoic acid and coenzyme A biosynthesis, and competence (**Fig. 8F and D**, **Table 2, Suppl. Table 4**). Genes showing increased expression in serum for both mutant strains included *ply* (encodes the extracellular toxin pneumolysin) and operons involved in fatty acid biosynthesis, purine biosynthesis, and two bacteriocin systems. To further characterise the metabolic responses of each mutant strain cultured in serum, the pathways enriched amongst the upregulated genes were identified using the KEGG database biological pathways annotations for the *S. pneumoniae* strain SP670-6B and over representation analysis (ORA) (57, 58). The Δ*proABC* showed significant enrichment for pathways involved in fatty acid biosynthesis, galactose metabolism, PTS systems, and amino acid and sugar metabolism (**Fig. 9A**). The Δ*fhs* strain had enriched expression of genes from multiple metabolic pathways including biosynthesis of secondary metabolites, competence, and purine, pyruvate, propanoate, amino acid and sugar metabolism (**Fig. 9B**).

**Figure 9.**
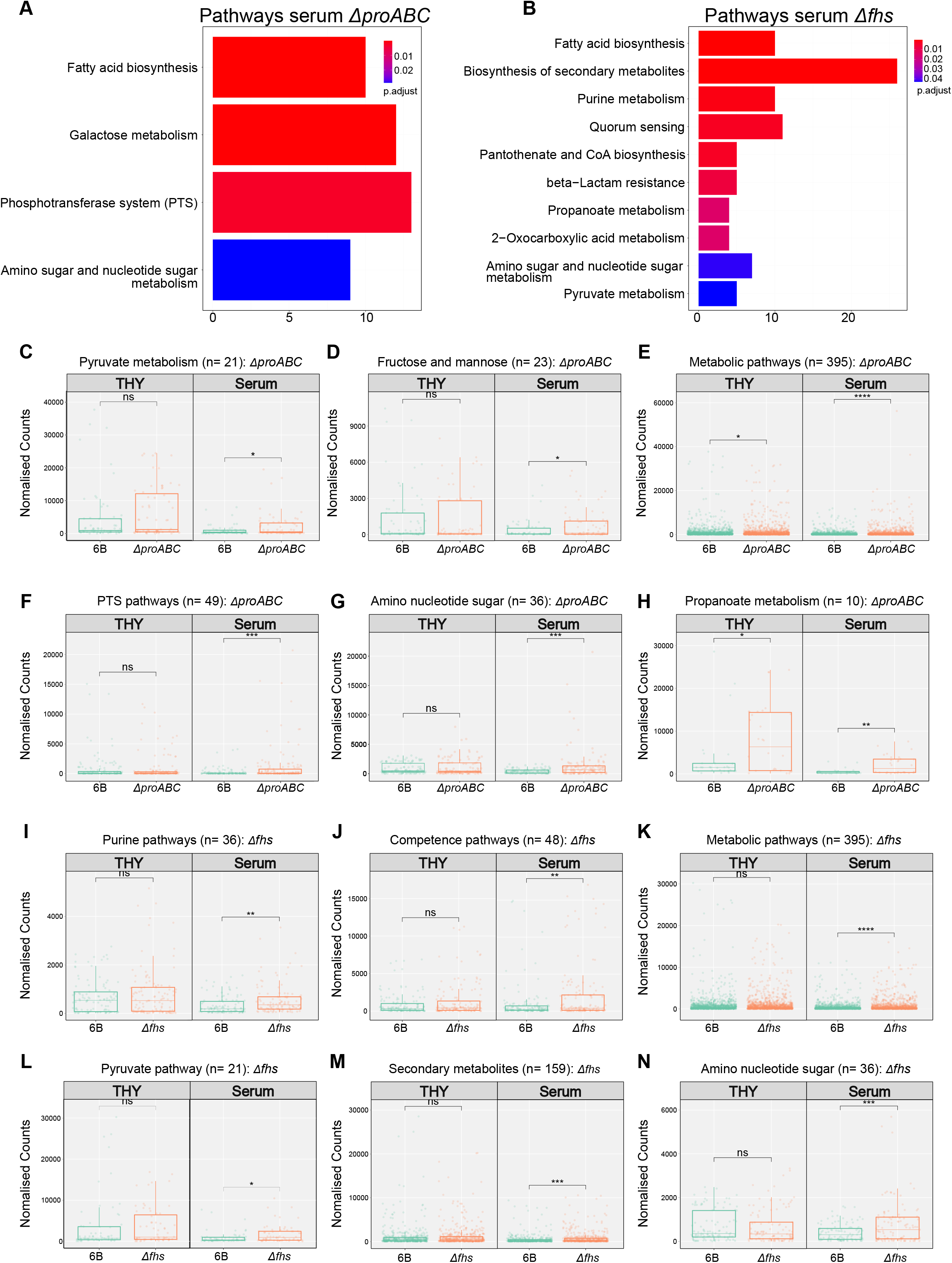
Pathway enrichment analysis for growth of mutants in THY or human serum, compared to wild-type. Pathways enriched amongst the upregulated genes within (**A**) Δ*proabc* and (**B**) Δ*fhs* mutant strains cultured in serum were identified using the KEGG database biological pathways annotations for the *S. pneumoniae* strain SP670-6B and ORA. (**C-N**) Overall expression of all genes within six selected metabolic pathways chosen from the pathways enrichment were compared for Δ*proABC* (**C-H**) and Δ*fhs* (**I-N**) strains using both the THY and serum RNAseq data.

**Table 2.**
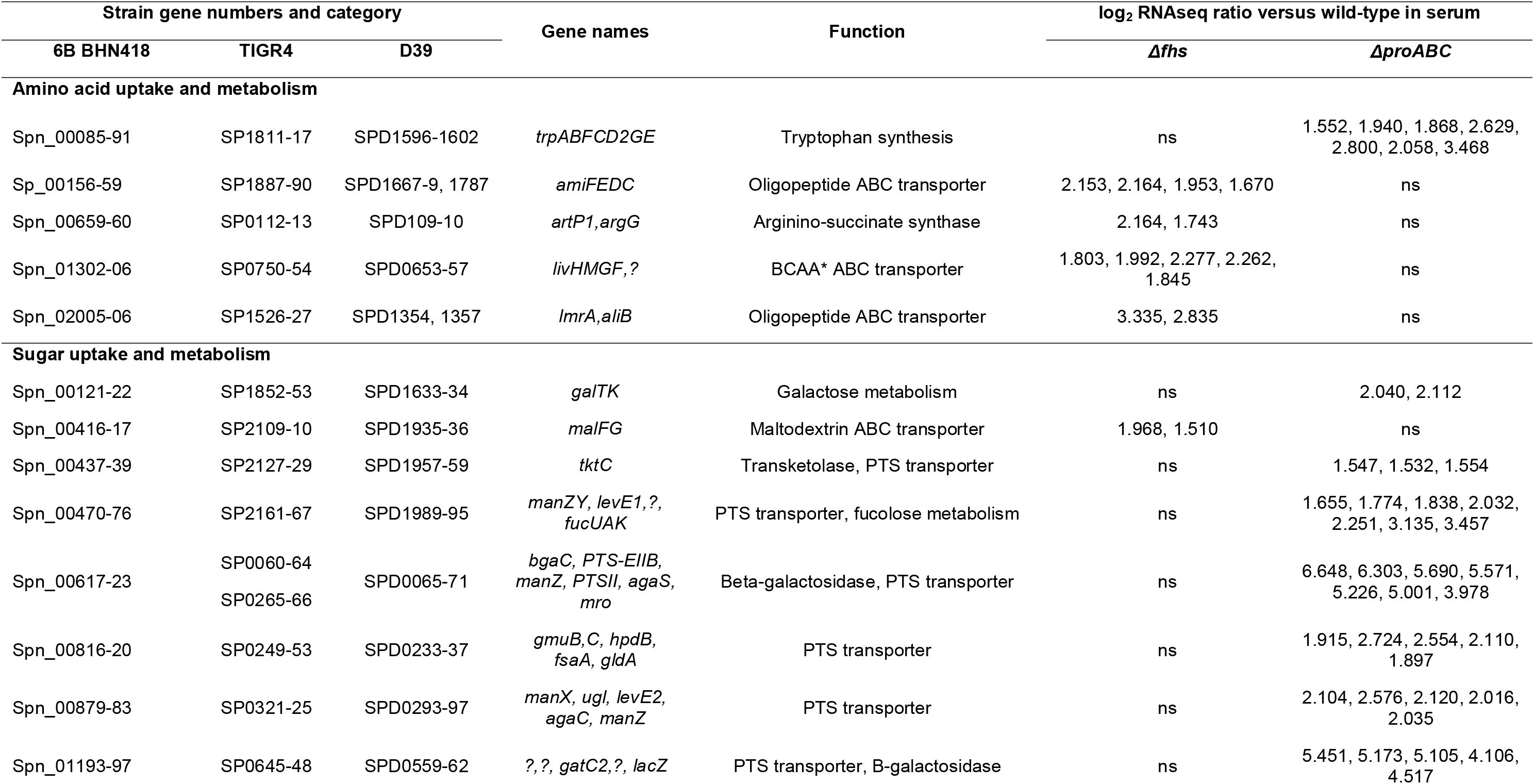
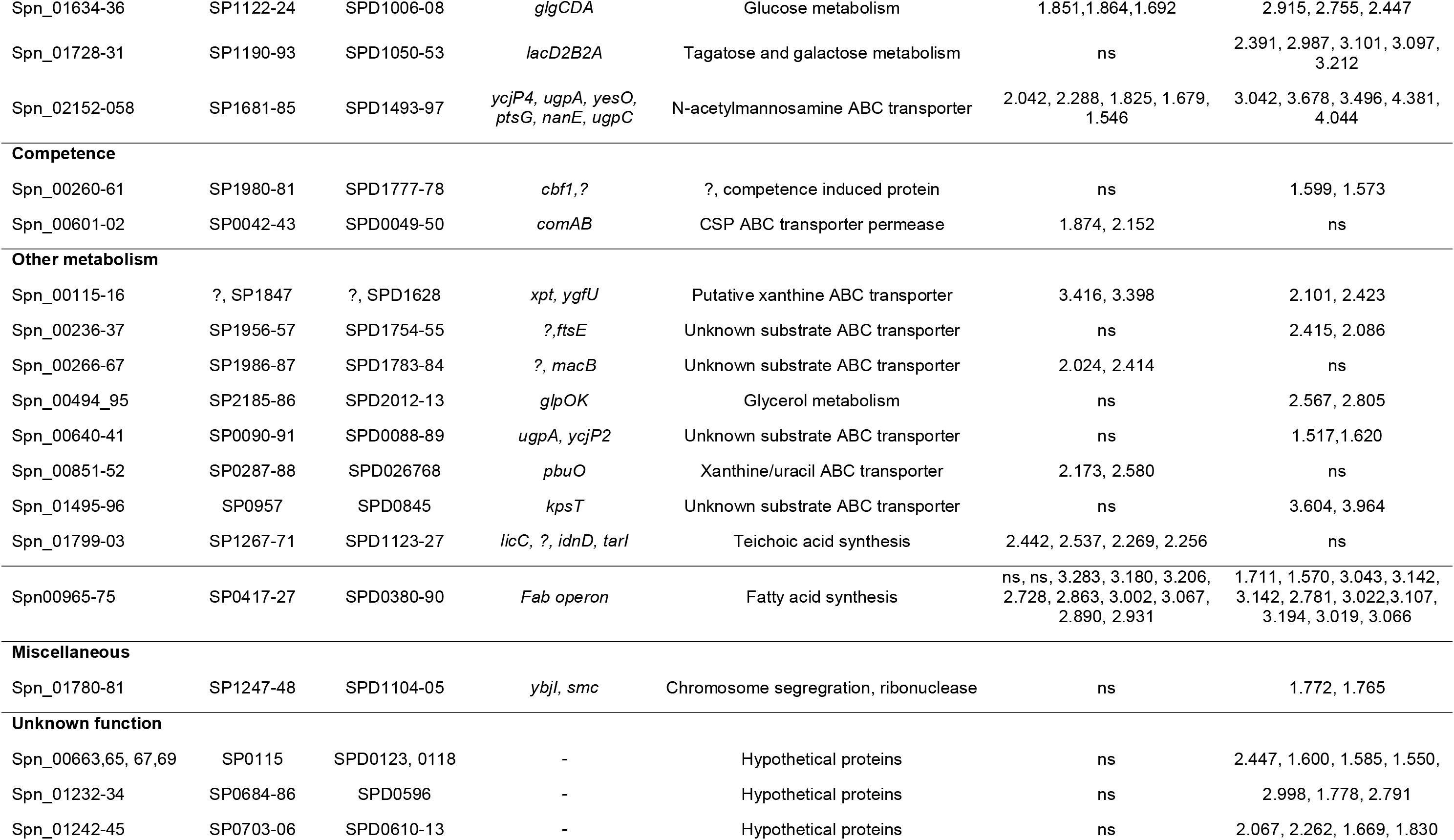
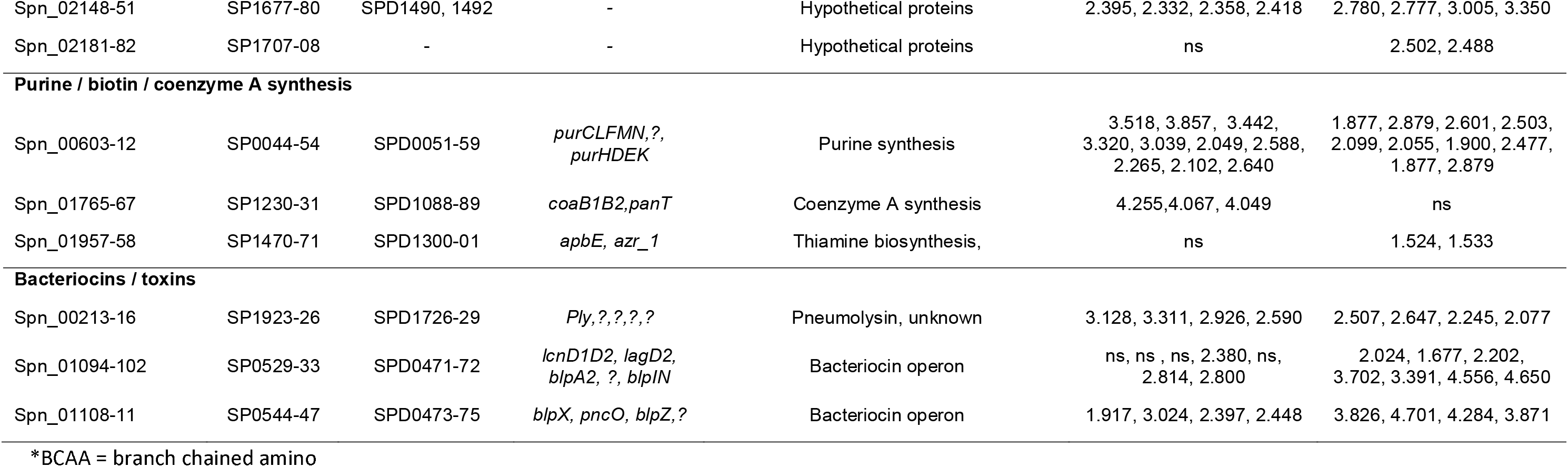
Gene operons showing differential expression (log_2_ fold change) between the mutant strains Δ*proABC* or Δ*fhs* and the wild-type 6B strain specifically when cultured in human serum (excluding those also showing differences in THY).

The above pathway analyses only use data for genes showing differential expression at a predetermined cut-off level. Hence, the overall expression of all genes within six selected metabolic pathways chosen from the above results for each strain were compared for each strain using both the THY and serum RNAseq data (**Fig. 9C to N**). When cultured in THY the Δ*proABC* and the Δ*fhs* strains showed significant increases in expression of genes from two (Fig. **9E** and **9H**) and none respectively of the six pathways assessed. In contrast, both mutant strains showed significant increases in gene expression compared to wild type for all six pathways when cultured in serum. These results further demonstrated that compared to complete media, culture in serum triggered multiple compensatory metabolic responses by the Δ*proABC* and the Δ*fhs* strains. Collectively the transcriptome data demonstrated that for *S. pneumoniae* cultured in human sera, loss of *fhs* or *proABC* increased transcription of multiple genes largely representing different metabolic processes for each strain, reflecting the specific effects of loss of *fhs* or the *proABC* operon on *S. pneumoniae* physiology during systemic infection.

## DISCUSSION

Using deletion mutants we have investigated the roles of the *S. pneumoniae* genetic loci *fhs* and *proABC* which are predicted to be important for different key aspects of bacterial metabolism. Both the Δ*fhs* and Δ*proABC S. pneumoniae* BHN418 capsular serotype 6B strains were severely attenuated in virulence in mouse models, with similar levels of recovery of target organ CFU as the unencapsulated mutant and 90%+ survival in the pneumonia model. Further characterisation *in vitro* demonstrated that the reduced virulence of the 6B Δ*fhs* and Δ*proABC* strains was likely to be related to poor growth in serum or CSF, phenotypes that will largely prevent *S. pneumoniae* from causing septicaemia or meningitis respectively. Culture under specific stress conditions identified significant differences between the mutant strains, with the Δ*proABC* showing increased sensitivity to osmotic or oxidative stress but not the Δ*fhs* strain. These results were supported by RNAseq data which demonstrated major differences in the Δ*proABC* and Δ*fhs* strain transcriptional response to culture in serum, representing activation of different metabolic pathways in response to the effects of each mutation on growth under infection conditions. Overall, these data demonstrate that both Fhs and ProABC have important roles for *S. pneumoniae* virulence related to their central importance for different metabolic pathways that are required for bacterial adaptation to growth in serum or CSF.

The amino acid proline is critical for protein synthesis, and can either be synthesized from glutamate or acquired from the environment via proline transporters (59, 60). Our data demonstrated that the BHN418 6B Δ*proABC* mutant strain was unable to grow in CDM, which contains 0.1mg/ml of proline. Supplementation with 1mg/ml proline restored partial growth in CDM demonstrating that the growth defect of the Δ*proABC* mutant was linked to loss of proline synthesis. These results demonstrate a central role for proline synthesis from glutamate by ProABC for *S. pneumoniae* growth, loss of which can not be overcome by uptake when there are only relatively low levels of environmental proline. The *S. pneumoniae* proline transporters have yet to be described, with no know equivalent transporters to the high affinity proline transporters of *Bacillus subtilis* (*opuE*) (59, 60) or *S. aureus* (*putP* and *proP*) (61). Concentrations of proline in human serum (0.002 mg/ml) are far lower than in CDM (62), explaining why the 6B Δ*proABC* mutant was unable to grow in serum or CSF without proline supplementation. However, proline supplementation with 0.1 mg/ml that was inadequate for growth in CDM partially restored growth of the 6B Δ*proABC* mutant strain in serum and CSF. Why a lower concentration of proline can restore growth in serum or CSF but not CDM is not clear; possibly serum and CSF have relatively higher proline availability from peptides or proteins (although supplementation with proline containing oligopeptides failed to complement our Δ*proABC* mutant), or higher concentrations of other nutrients that can compensate for the physiological defects caused by loss of proline synthesis. We used RNAseq to characterise how loss of proline synthesis affects *S. pneumoniae* metabolism during growth, reasoning that genes showing increased expression by the Δ*proABC* mutant in serum compared to the wild-type strain will represent compensatory metabolic pathways activated in response to loss of proline availability. Perhaps surprisingly, the differences in gene expression by the Δ*proABC* mutant compared to the wild type when cultured in serum were dominated by increased expression of pathways related to carbohydrate uptake and metabolism rather than amino acids. This could indicate that proline is an important alternative carbon source for *S. pneumoniae* growth in serum or CSF. As well as its role for biosynthesis, proline availability could also affect *S. pneumoniae* growth in serum through its important role for osmoregulation (63, 64). High cytoplasmc concentrations of proline can protect bacteria against osmotic stress without affecting other cellular functions (29, 64), and proline transport protects against osmotic stress and improves virulence for *E. coli* (65) and *S. aureus* (66). Our data confirmed proline is likely to be important for osmoregulation of *S. pneumoniae* as well, with the Δ*proABC* mutant showing increased sensitivity to osmotic stress, but unlike *S. aureus* (the genome of which does not contain *proA* or *proB*) this is dependent on proline synthesis rather than uptake (67). In addition, loss of proline synthesis could potentially affect *S. pneumoniae* virulence through effects on the synthesis of virulence proteins such as PspC and PspA which contain proline-rich domains (68, 69).

Although *fhs* was identified by separate *S. pneumoniae* virulence screens (24, 26) and is required for *Streptococcus suis* virulence (70), the role of Fhs for bacterial virulence seems to have been largely under-appreciated. We found that *S. pneumoniae* growth in serum or CSF and therefore systemic virulence was totally dependent on *fhs*, demonstrating a central role of one carbon metabolism and probably the stringent stress response (37, 41) for *S. pneumoniae* metabolism under physiological conditions. Various studies have identified specifc metabolic roles for Fhs in other bacterial species, including anaerobic growth for *E. coli* (39), purine synthesis for *Streptococcus mutans* (36), and folate homeostasis for *Clostridia perfringens* (41). We found that although addition of exogenous purines did not restore growth of the *S. pneumoniae* Δ*fhs* strain in CDM, it did partially restore growth in serum. Furthermore, the RNAseq data demonstrated upregulation of purine pathways when the *S. pneumoniae* Δ*fhs* strain was cultured in serum. These data suggest *S. pneumoniae* purine metabolism is dependent on Fhs, and provide further support that serum is more permissive for *S. pneumoniae* growth than CDM as suggested by the data described above for the Δ*proABC* strain. The RNAseq data also showed loss of *fhs* increased expression of a broad range of metabolic pathways, compatible with the role of the one carbon metaboloism in the synthesis of multiple products including amino acids. The increased expression of lipid metabolism and beta lactam resistance genes by the Δ*fhs* strain in serum suggest an important role for Fhs and the one carbon pathway in maintaining the cell envelope. In addition, the RNAseq showed deletion of *fhs* increased expression of quorum sensing pathways, perhaps as a component of the *S. pneumoniae* response to physiological stress.

Depsite the severe attenuation of the 6B Δ*fhs* and Δ*proABC* strains for systemic virulence these strains were still able to persist in the nasopharynx and establish significant levels of lung CFU, a difference we have previously shown can be exploited to make live-attenuated *S. pneumoniae* vaccines (42, 43). Why the physiological conditions in the respiratory tract are less dependent on proline synthesis and the one carbon metabolism pathways than in serum is not clear. Glucose is the main carbohydrate source in blood or CSF, whereas in the nasopharynx glucose is only present at very low levels and several alternative carbohydates such as galactose are available to support *S. pneumoniae* growth (71). Growth of the Δ*proABC* strain was similar to the wild type strain in the artificial nose-like media containing N-Acetylglucosamine, suggesting the available carbon source may cause the differential survival of this mutant strain during nasopharyngeal colonisation compared to systemic infection. Furthermore, during mouse infection wild type *S. pneumoniae* CFU increased in the blood from zero to approximately 10^4^ within 24 hours, whereas in the nasopharynx or lung recovered CFU/ml were lower than the initial inoculum. The rapid growth in serum could increase the importance of specific metabolic pathways supported by ProABC and Fhs for bacterial survival compared to the slower replicatipn in the nasopharynx.

*S. pneumoniae* essential genes can be divided into universal, core-strain specific (present in all strains but not always essential), and accessory essential genes (essential but only present in some strains) categories (37). *fhs* has been described as a core-strain specific essential gene, although we and others have readily obtained Δ*fhs* strains in several *S. pneumoniae* strain backgrounds (24, 26) using blood agar as the culture medium. However, *fhs* was essential for growth in blood, CSF, nose- mimicking media, or CDM illustrating that which genes are essential is highly dependent on the *in vitro* and *in vivo* growth conditions; we would argue that for bacterial pathogens gene essentiality should be based on the physiological conditions found at their main site(s) of infection. Transfer of the Δ*proABC* mutations to the D39 *S. pneumoniae* strain background demonstrated that unlike the 6B Δ*proABC* strain the D39 Δ*proABC* strain was able to cause septicaemia in the sepsis model and replicate in blood *ex vivo*, albeit at a reduced level to the wild-type strain. These results show that the essential role of *S. pneumoniae* ProABC for replication in blood and therefore virulence is strain- dependent. Hence, similar to essential genes, bacterial virulence genes can be divided into universal and core-strain specific categories reflecting differences between strains in growth requirements under physiological conditions. These differences in metabolic function could underpin some of the differences seen between *S. pneumoniae* strains in virulence, and similar mechanisms are also likely to be relevant for multiple other bacterial pathogens.

In conclusion, we have demonstrated that Fhs and and therefore one carbon metabolism has an important role for *S. pneumoniae* virulence related to its central importance for multiple metabolic pathways required for growth in human serum and CSF. The one carbon metabolism pathway is present in many bacterial species and hence our data could be relevant for multiple other bacterial pathogens. In addition, we have identified a strain-dependent role for proline biosynthesis for *S. pneumoniae* growth in serum and for virulence, demonstrating metabolic functions can influence differential virulence between different *S. pneumoniae* strains and perhaps for other pathogenic bacteria. These results emphasise the importance of factors controlling bacterial metabolism for the development of infection, and form the basis for further work characterising in greater detail the metabolic pathways needed for bacterial virulence.

## METHODS

### Bacterial strains and growth conditions

Bacteria were grown in Todd-Hewitt broth (THY) (Sigma) supplemented with 0.5% yeast extract (Sigma) and cultured in the presence of 5% CO_2_ at 37°C or in Columbia agar supplemented with 5% horse blood (Oxoid) in solid media. Bacteria were stored as single aliquots of 0.5 ml in THY broth at - 80°C with 15% glycerol (O.D_595_ 0.4-0.5), thawed and diluted in PBS to appropriate concentration as required. Plasmids and mutants strains were selected using spectinomycin (Spec) 150 μg/ml or kanamycin (Kan) 250 μg/ml. Replication of *S. pneumoniae* strains in THY, chemically defined medium (CDM), human sera or cerebrospinal fluid (CSF) from healthy volunteers, was determined by inoculating 5×10^6^ CFU/well in a 200μl volume and measuring change in OD_595_ every 30 minutes using a TECAN Spark® plate reader. Stress conditions were generated as necessary by addition of: oxidative stress, up to 5 mM paraquat (Sigma-Aldrich); cation restriction, addition of ethylene diamine di-o-hydroxyphenylacetic (EDDA) at 200 μM; increased osmolarity, 50, 100, or 200 mM NaCl or 100 or 400 mM sucrose. When required CDM media or serum / CSF was supplemented with (i) different carbon sources at 1% (mannose, maltose, N-Acetyl glucosamine, glucose, or a mix of the four sugars 12.5mM each), (ii) proline 0.1 or 1 mg/ml proline, (iii) oligopeptides pro8x (PPPPPPPP), AliAPro (FNEMQPIVDRQPPPP) or AliBPro (AIQSEKARKHNPPPP) (Nasher 2018) at 10, 50 or 250 ug/ml, or (iv) purines, adenine and/or glycine at 1 mg/ml.

### Construction of Δ*cps*, Δ*proABC*, Δ*fhs and* Δ*fhs*+fhs *S. pneumoniae* strains

Plasmids and primers used in this study are described in **Suppl Table 1**. Mutant strains were constructed by overlap extension PCR as previously described (72) using a transformation fragment in which the gene of interest has been replaced for either Spec (Δ*proABC*, Δ*fhs*) or Kan cassette (Δ*cps*) amplified from plasmids pR412 and pBAG5 respectively (73, 74). Briefly, two products corresponding to 600-1000 bp 5′ and 3′ to the target gene were amplified from genomic DNA by PCR carrying 3′ and 5′ linkers complementary to the sequence of the antibiotic resistance gene, and the fragments fused by overlap extension PCR then transformed into *S. pneumoniae* by homologous recombination and allelic replacement using a mix of competence stimulating peptides (CSP-1 and CSP-2) (kind gift from D. Morrison) and antibiotic selection (75). Gene deletions were confirmed by PCR and sequencing of the PCR products. Mutation stability was confirmed by multiples rounds of growth in THY without antibiotics then plating into blood agar plates with and without antibiotics (**data not shown**). The Δ*fhs* strain was complemented by ectopic insertion of a copy of fhs using the promoterless integrative plasmid pPEPY, (kind gift from Jan-Willem Veening) (Addgene plasmid # 122633) (49).

### Mouse infection models

Mouse infection experimental procedures were approved by the local ethical review process and performed according to UK national guidelines under the UK Home Office project licence PPL70/6510. Groups of outbred CD1 female mice (Charles River Breeders) 4 to 6 weeks old were infected with *S. pneumoniae* by intraperitoneal (IP) injection (5×10^6^ CFU n 100 µl, sepsis model) of, or by intranasal (IN) inoculation under isoflurane anaesthesia for the pneumonia (1×10^7^ CFU bacteria in 50 µl) or nasopharyngeal colonisation (1×10^7^ CFU bacteria in 10 µl) models. Target organs (nasal washes, lung and spleen homogenates, or blood) were recovered at pre-specified time points and CFU concentrations calculated by plating serial dilutions onto blood agar plates and overnight culture (14, 76)

### Flow cytometry C3b, IgG and phosphocholine binding and neutrophil killing assays

Binding of complement component C3b/iC3b or IgG when incubated in human sera, to live *S. pneumoniae* were detected by flow cytometry as previously described (77) and the following secondary antibodies; fluorescein isothiocyanate-conjugated polyclonal goat anti-human C3b antibody (ICN) and PE-conjugated goat anti-mouse IgG (Biolegend). For the neutrophil killing assays, *S. pneumoniae* strains previously incubated in 25% of baby rabbit complement (BioRad) at 37°C for 30 mins were added to fresh human neutrophils extracted from blood from healthy volunteers using MACSxpress kit (Miltenyi Biotec) at an MOI of 1:100 (72) before plating serial dilutions of supernatants onto blood agar plates to calculate surving CFU. The results expressed as a percentage of CFU recovered relative to wild-type strain.

### Serum and CSF sources

Human serum from healthy volunteers unvaccinated against *S. pneumoniae* was obtained after obtaining informed consent according to institutional guidelines and stored as single-use aliquots at −80°C as a source of complement and serum components. CSF kind gift from Diederik van de Beek at UMC, Netherlands.

### RNA methods

Triplicate cultures of each *S. pneumoniae* strains were grown in THY broth until an OD_595_ 0.4-0.5, then centrifuged, resuspended in RNAprotect stabilization buffer (Qiagen), centrifuged again before RNA extraction using Mirvana RNA kit (Applied biosystems) according to the manufacturer’s instructions with an additional lysis step using vigoruous shaking of the tubes after addition of 0.1 mm glass beads (MP Biomedicals). For the serum studies, bacteria were grown in THY broth up to OD_595_ 0.4-0.5, washed twice in PBS and resuspended in 500 ul of fresh human sera for 60 minutes following the same protocol mentioned above before RNA extraction. Purified RNA was treated with Turbo DNAse (Applied biosystems), ribosomal RNA removed using MICROBExpress™ (Thermo scientific), and 100 ng used to construct libraries for sequencing using the KAPA RNA HyperPrep kit (Roche Diagnostics) with 8 cycles of amplification. Qubit (Thermo Fisher Scientific) was used to assess library concentrations and TapeStation (Agilent Technologies) confirmed library sizes. Libraries were multiplexed to 24 samples per run and single-end sequenced by the UCL Pathogen Genomics Unit (PGU) with the NextSeq 500 desktop sequencer (Illumina) using a 75 cycle high-output kit. The quality of raw FASTQ reads was checked by FastQC v0.11.5, Babraham Bioinformatics, UK (78) and visualised using multiQC v1.9(79). Reads were trimmed using Trimmomatic v0.39 (80). Trimmed reads were also checked by FastQC and multiQC. Trimmed reads were then mapped to the KEGG annotated *S. pneumoniae* serotype 6B genome sequence (670-6B, Accession: CP002176.1) using bowtie2 v2.4.4 with default settings (81). Conversion into BAM files was performed using SAMtools (82). Mapped reads were visualized in the Integrated Genome Browser (IGB) (83). FeatureCounts v2.0.0 was used to summarize read counts for each annotated feature in multimapping mode (-M) (84).

### Differential gene expression analyses

The generated count matrix was imported into R-studio (R v3.4.2) and normalization of counts and differential gene expression analysis was performed by DESeq2 (85). DESeq2 normalized libraries were regularized log transformed for visualizing heatmaps and clustering. Differential gene expression was performed on raw counts. Features (genes) with a log2 fold change > 1.5 and false discovery rate (FDR) of < 0.05 were categorized as differentially expressed. KEGG pathway enrichment and module analysis were performed in R studio using clusterProfiler (86).

### Statistical analysis

Statistical analyses were performed using GraphPad Prism 8 (GraphPad Software, La Jolla, CA, USA) or R. (R v3.4.2). Quantitative results are expressed as median and interquartile range for animal experiments, and analsyed using the Kruskal-Wallis non-parametric test. Dunn’s multiple comparisons test were used for *post hoc* analysis. *P*-values < 0.05 (95% confidence) were considered statistically significant.

## Data availability

Raw RNA-seq data have been deposited in the ArrayExpress database at EMBL-EBI (www.ebi.ac.uk/arrayexpress,) under accession number XXXX

## Acknowledgements

This work was supported by MRC grants R/N02687X/1 and MR/R001871/1, and undertaken at UCLH/UCL who receive funding from the Department of Health’s NIHR Biomedical Research Centre’s funding scheme. The authors would like to thank the Pathogens Genomic Unit, an initiative established by grants from the Medical Research Council and the UCL/UCLH, for carrying the RNA sequencing.

## Supplementary figures

**Supplementary figure 1. Effect of proline-containing peptides on** Δ***proABC* mutant growth** Growth of wild-type and Δ*proABC* mutant strain in CDM complemented by (**A**) pro8x peptide (PPPPPPPP), (**B**) AliA (FNEMQPIVDRQPPPP) or (**C**) AliB (AIQSEKARKHNPPPP) at three different concentrtions; 250, 50 and 10 ug ml^-1^. Differences in growth were assessed by measuring OD_595_ every 30 min for a 24 hours period.

**Supplementary figure 2.** Principal component analysis of normalised and transformed transcripts for 6B, Δ*proABC* and Δ*fhs* strains in human serum and THY.

## References

1. O’Brien KL, Wolfson LJ, Watt JP, Henkle E, Deloria-Knoll M, McCall N, Lee E, Mulholland K, Levine OS, Cherian T, Hib, Pneumococcal Global Burden of Disease Study T. 2009. Burden of disease caused by Streptococcus pneumoniae in children younger than 5 years: global estimates. Lancet 374:893–902.

2. Melegaro A, Edmunds WJ, Pebody R, Miller E, George R. 2006. The current burden of pneumococcal disease in England and Wales. J Infect 52:37–48.

3. Goldblatt D, Hussain M, Andrews N, Ashton L, Virta C, Melegaro A, Pebody R, George R, Soininen A, Edmunds J, Gay N, Kayhty H, Miller E. 2005. Antibody responses to nasopharyngeal carriage of Streptococcus pneumoniae in adults: a longitudinal household study. J Infect Dis 192:387–93.

4. Brooks LRK, Mias GI. 2018. Streptococcus pneumoniae’s virulence and host immunity: aging, diagnostics, and prevention. Front Immunol 9:1366.

5. Hyams C, Camberlein E, Cohen JM, Bax K, Brown JS. 2010. The Streptococcus pneumoniae capsule inhibits complement activity and neutrophil phagocytosis by multiple mechanisms. Infect Immun 78:704–15.

6. Hyams C, Trzcinski K, Camberlein E, Weinberger DM, Chimalapati S, Noursadeghi M, Lipsitch M, Brown JS. 2013. Streptococcus pneumoniae capsular serotype invasiveness correlates with the degree of factor H binding and opsonization with C3b/iC3b. Infect Immun 81:354–63.

7. Pracht D, Elm C, Gerber J, Bergmann S, Rohde M, Seiler M, Kim KS, Jenkinson HF, Nau R, Hammerschmidt S. 2005. PavA of Streptococcus pneumoniae modulates adherence, invasion, and meningeal inflammation. Infect Immun 73:2680–9.

8. Barocchi MA, Ries J, Zogaj X, Hemsley C, Albiger B, Kanth A, Dahlberg S, Fernebro J, Moschioni M, Masignani V, Hultenby K, Taddei AR, Beiter K, Wartha F, von Euler A, Covacci A, Holden DW, Normark S, Rappuoli R, Henriques-Normark B. 2006. A pneumococcal pilus influences virulence and host inflammatory responses. Proc Natl Acad Sci U S A 103:2857–62.

9. Quin LR, Onwubiko C, Moore QC, Mills MF, McDaniel LS, Carmicle S. 2007. Factor H binding to PspC of Streptococcus pneumoniae increases adherence to human cell lines in vitro and enhances invasion of mouse lungs in vivo. Infect Immun 75:4082–7.

10. Camberlein E, Cohen JM, Jose R, Hyams CJ, Callard R, Chimalapati S, Yuste J, Edwards LA, Marshall H, van Rooijen N, Noursadeghi M, Brown JS. 2015. Importance of bacterial replication and alveolar macrophage-independent clearance mechanisms during early lung infection with Streptococcus pneumoniae. Infect Immun 83:1181–9.

11. Marshall H, Jose RJ, Kilian M, Petersen FC, Brown JS. 2021. Effects of expression of Streptococcus pneumoniae PspC on the ability of Streptococcus mitis to evade complement-mediated immunity. Front Microbiol 12:773877.

12. Weiser JN, Ferreira DM, Paton JC. 2018. Streptococcus pneumoniae: transmission, colonization and invasion. Nat Rev Microbiol 16:355–367.

13. Man WH, de Steenhuijsen Piters WA, Bogaert D. 2017. The microbiota of the respiratory tract: gatekeeper to respiratory health. Nat Rev Microbiol 15:259–270.

14. Brown JS, Gilliland SM, Holden DW. 2001. A Streptococcus pneumoniae pathogenicity island encoding an ABC transporter involved in iron uptake and virulence. Mol Microbiol 40:572–85.

15. Brown JS, Gilliland SM, Ruiz-Albert J, Holden DW. 2002. Characterization of pit, a Streptococcus pneumoniae iron uptake ABC transporter. Infect Immun 70:4389–98.

16. Novak R, Cauwels A, Charpentier E, Tuomanen E. 1999. Identification of a Streptococcus pneumoniae gene locus encoding proteins of an ABC phosphate transporter and a two-component regulatory system. J Bacteriol 181:1126–33.

17. Orihuela CJ, Mills J, Robb CW, Wilson CJ, Watson DA, Niesel DW. 2001. Streptococcus pneumoniae PstS production is phosphate responsive and enhanced during growth in the murine peritoneal cavity. Infect Immun 69:7565–71.

18. Ware D, Jiang Y, Lin W, Swiatlo E. 2006. Involvement of potD in Streptococcus pneumoniae polyamine transport and pathogenesis. Infect Immun 74:352–61.

19. Bayle L, Chimalapati S, Schoehn G, Brown J, Vernet T, Durmort C. 2011. Zinc uptake by Streptococcus pneumoniae depends on both AdcA and AdcAII and is essential for normal bacterial morphology and virulence. Mol Microbiol 82:904–16.

20. Brown JS, Gilliland SM, Basavanna S, Holden DW. 2004. phgABC, a three-gene operon required for growth of Streptococcus pneumoniae in hyperosmotic medium and in vivo. Infect Immun 72:4579–88.

21. Chimalapati S, Cohen J, Camberlein E, Durmort C, Baxendale H, de Vogel C, van Belkum A, Brown JS. 2011. Infection with conditionally virulent Streptococcus pneumoniae Δpab strains induces antibody to conserved protein antigens but does not protect against systemic infection with heterologous strains. Infect Immun 79:4965–76.

22. Hava DL, Camilli A. 2002. Large-scale identification of serotype 4 Streptococcus pneumoniae virulence factors. Mol Microbiol 45:1389–406.

23. Johnson MD, Echlin H, Dao TH, Rosch JW. 2015. Characterization of NAD salvage pathways and their role in virulence in Streptococcus pneumoniae. Microbiology 161:2127–36.

24. van Opijnen T, Camilli A. 2012. A fine scale phenotype-genotype virulence map of a bacterial pathogen. Genome Res 22:2541–51.

25. Ogunniyi AD, Mahdi LK, Trappetti C, Verhoeven N, Mermans D, Van der Hoek MB, Plumptre CD, Paton JC. 2012. Identification of genes that contribute to the pathogenesis of invasive pneumococcal disease by in vivo transcriptomic analysis. Infect Immun 80:3268–78.

26. Mahdi LK, Wang H, Van der Hoek MB, Paton JC, Ogunniyi AD. 2012. Identification of a novel pneumococcal vaccine antigen preferentially expressed during meningitis in mice. J Clin Invest 122:2208–20.

27. Belitsky BR, Brill J, Bremer E, Sonenshein AL. 2001. Multiple genes for the last step of proline biosynthesis in Bacillus subtilis. J Bacteriol 183:4389–92.

28. Bremer E. 2002. Adaptation to changing osmolarity. In Bacillus subtilis and its closest relatives: from genes to cells, p 385–391. In Sonenshein AL, Hoch, J.A., Losick, R. (ed). ASM Press, Washington, DC.

29. Kempf B, Bremer E. 1998. Uptake and synthesis of compatible solutes as microbial stress responses to high-osmolality environments. Arch Microbiol 170:319–30.

30. Brill J, Hoffmann T, Bleisteiner M, Bremer E. 2011. Osmotically controlled synthesis of the compatible solute proline is critical for cellular defense of Bacillus subtilis against high osmolarity. J Bacteriol 193:5335–46.

31. Christgen SL, Becker DF. 2019. Role of proline in pathogen and host interactions. Antioxid Redox Signal 30:683–709.

32. Smith DA, Parish T, Stoker NG, Bancroft GJ. 2001. Characterization of auxotrophic mutants of Mycobacterium tuberculosis and their potential as vaccine candidates. Infect Immun 69:1142–50.

33. Smith AP, Lane LC, van Opijnen T, Woolard S, Carter R, Iverson A, Burnham C, Vogel P, Roeber D, Hochu G, Johnson MDL, McCullers JA, Rosch J, Smith AM. 2021. Dynamic pneumococcal genetic adaptations support bacterial growth and inflammation during coinfection with influenza. Infect Immun 89:e0002321.

34. Molzen TE, Burghout P, Bootsma HJ, Brandt CT, van der Gaast-de Jongh CE, Eleveld MJ, Verbeek MM, Frimodt-Moller N, Ostergaard C, Hermans PW. 2011. Genome-wide identification of Streptococcus pneumoniae genes essential for bacterial replication during experimental meningitis. Infect Immun 79:288–97.

35. Afzal M, Shafeeq S, Kuipers OP. 2016. Methionine-mediated gene expression and characterization of the CmhR regulon in Streptococcus pneumoniae. Microb Genom 2:e000091.

36. Crowley PJ, Gutierrez JA, Hillman JD, Bleiweis AS. 1997. Genetic and physiologic analysis of a formyl-tetrahydrofolate synthetase mutant of Streptococcus mutans. J Bacteriol 179:1563–72.

37. Rosconi F, Rudmann E, Li J, Surujon D, Anthony J, Frank M, Jones DS, Rock C, Rosch JW, Johnston CD, van Opijnen T. 2022. A bacterial pan-genome makes gene essentiality strain-dependent and evolvable. Nat Microbiol 7:1580–1592.

38. Paukert JL, Rabinowitz JC. 1980. Formyl-methenyl-methylenetetrahydrofolate synthetase (combined): a multifunctional protein in eukaryotic folate metabolism. Methods Enzymol 66:616–26.

39. Sah S, Aluri S, Rex K, Varshney U. 2015. One-carbon metabolic pathway rewiring in Escherichia coli reveals an evolutionary advantage of 10-formyltetrahydrofolate synthetase (Fhs) in survival under hypoxia. J Bacteriol 197:717–26.

40. Whitehead TR, Park M, Rabinowitz JC. 1988. Distribution of 10-formyltetrahydrofolate synthetase in eubacteria. J Bacteriol 170:995–7.

41. Aluri S, Sah S, Miryala S, Varshney U. 2016. Physiological role of FolD (methylenetetrahydrofolate dehydrogenase), FchA (methenyltetrahydrofolate cyclohydrolase) and Fhs (formyltetrahydrofolate synthetase) from Clostridium perfringens in a heterologous model of Escherichia coli. Microbiology 162:145–155.

42. Ramos-Sevillano E, Ercoli G, Felgner P, Ramiro de Assis R, Nakajima R, Goldblatt D, Heyderman RS, Gordon SB, Ferreira DM, Brown JS. 2020. Preclinical development of virulence attenuated Streptococcus pneumoniae strains able to enhance protective immunity against pneumococcal infection. Am J Respir Crit Care Med doi:10.1164/rccm.202011-4161LE.

43. Hill H, Mitsi E, Nikolaou E, Blizard A, Pojar S, Howard A, Hyder-Wright A, Devin J, Reiné J, Robinson R, Solórzano C, Jochems S, Kenny-Nyazika T, Ramos-Sevillano E, Weight CM, Myerscough C, McLennan D, Morton B, Gibbons E, Farrar M, Randles V, Burhan H, Chen T, Shandling AD, Campo JJ, Heyderman R, Gordon SB, Brown JS, Collins AM, Ferreira DM. 2023. A randomised controlled trial of nasal immunisation with live virulence attenuated Streptococcus pneumoniae strains using human infection challenge. Am J Respir Crit Care Med. In press.

44. Csonka LN, Leisinger T. 2007. Biosynthesis of proline. EcoSal Plus 2.

45. Celeste LR, Chai G, Bielak M, Minor W, Lovelace LL, Lebioda L. 2012. Mechanism of N10- formyltetrahydrofolate synthetase derived from complexes with intermediates and inhibitors. Protein Sci 21:219–28.

46. Lovell CR, Przybyla A, Ljungdahl LG. 1990. Primary structure of the thermostable formyltetrahydrofolate synthetase from Clostridium thermoaceticum. Biochemistry 29:5687–94.

47. Cook RJ, Lloyd RS, Wagner C. 1991. Isolation and characterization of cDNA clones for rat liver 10-formyltetrahydrofolate dehydrogenase. J Biol Chem 266:4965–73.

48. Jones DT. 1999. Protein secondary structure prediction based on position-specific scoring matrices. J Mol Biol 292:195–202.

49. Keller LE, Rueff AS, Kurushima J, Veening JW. 2019. Three new integration vectors and fluorescent proteins for use in the opportunistic human pathogen Streptococcus pneumoniae. Genes (Basel) 10.

50. Bolm M, Jansen WT, Schnabel R, Chhatwal GS. 2004. Hydrogen peroxide-mediated killing of Caenorhabditis elegans: a common feature of different streptococcal species. Infect Immun 72:1192–4.

51. Garsin DA, Sifri CD, Mylonakis E, Qin X, Singh KV, Murray BE, Calderwood SB, Ausubel FM. 2001. A simple model host for identifying Gram-positive virulence factors. Proc Natl Acad Sci U S A 98:10892–7.

52. Csonka LN. 1989. Physiological and genetic responses of bacteria to osmotic stress. Microbiol Rev 53:121–47.

53. Hassett DJ, Britigan BE, Svendsen T, Rosen GM, Cohen MS. 1987. Bacteria form intracellular free radicals in response to paraquat and streptonigrin. Demonstration of the potency of hydroxyl radical. J Biol Chem 262:13404–8.

54. Nasher F, Aguilar F, Aebi S, Hermans PWM, Heller M, Hathaway LJ. 2018. Peptide ligands of AmiA, AliA, and AliB proteins determine pneumococcal phenotype. Front Microbiol 9:3013.

55. Aprianto R, Slager J, Holsappel S, Veening JW. 2018. High-resolution analysis of the pneumococcal transcriptome under a wide range of infection-relevant conditions. Nucleic Acids Res 46:9990–10006.

56. Bidossi A, Mulas L, Decorosi F, Colomba L, Ricci S, Pozzi G, Deutscher J, Viti C, Oggioni MR. 2012. A functional genomics approach to establish the complement of carbohydrate transporters in Streptococcus pneumoniae. PLoS One 7:e33320.

57. Kanehisa M. 2000. KEGG: Kyoto Encyclopedia of genes and genomes. Nucleic Acids Research 28:27–30.

58. Kanehisa M, Goto S. 2000. KEGG: kyoto encyclopedia of genes and genomes. Nucleic Acids Res 28:27–30.

59. Spiegelhalter F, Bremer E. 1998. Osmoregulation of the opuE proline transport gene from Bacillus subtilis: contributions of the sigma A- and sigma B-dependent stress-responsive promoters. Mol Microbiol 29:285–96.

60. von Blohn C, Kempf B, Kappes RM, Bremer E. 1997. Osmostress response in Bacillus subtilis: characterization of a proline uptake system (OpuE) regulated by high osmolarity and the alternative transcription factor sigma B. Mol Microbiol 25:175–87.

61. Schwan WR, Wetzel KJ. 2016. Osmolyte transport in Staphylococcus aureus and the role in pathogenesis. World J Clin Infect Dis 6:22–27.

62. Zhu K, Zhang S, Yue K, Zuo Y, Niu Y, Wu Q, Pan W. 2022. Rapid and nondestructive detection of proline in serum using near-infrared spectroscopy and partial least squares. J Anal Methods Chem 2022:4610140.

63. Csonka LN, Hanson AD. 1991. Prokaryotic osmoregulation: genetics and physiology. Annu Rev Microbiol 45:569–606.

64. Wood JM. 2011. Bacterial osmoregulation: a paradigm for the study of cellular homeostasis. Annu Rev Microbiol 65:215–38.

65. Wood JM. 2006. Osmosensing by bacteria. Sci STKE 2006:pe43.

66. Wetzel KJ, Bjorge D, Schwan WR. 2011. Mutational and transcriptional analyses of the Staphylococcus aureus low-affinity proline transporter OpuD during in vitro growth and infection of murine tissues. FEMS Immunol Med Microbiol 61:346–55.

67. Schwan WR, Coulter SN, Ng EY, Langhorne MH, Ritchie HD, Brody LL, Westbrock-Wadman S, Bayer AS, Folger KR, Stover CK. 1998. Identification and characterization of the PutP proline permease that contributes to in vivo survival of Staphylococcus aureus in animal models. Infect Immun 66:567–72.

68. Daniels CC, Coan P, King J, Hale J, Benton KA, Briles DE, Hollingshead SK. 2010. The proline-rich region of pneumococcal surface proteins A and C contains surface-accessible epitopes common to all pneumococci and elicits antibody-mediated protection against sepsis. Infect Immun 78:2163–72.

69. Melin M, Coan P, Hollingshead S. 2012. Development of cross-reactive antibodies to the proline-rich region of pneumococcal surface protein A in children. Vaccine 30:7157–60.

70. Zheng C, Xu J, Shi G, Zhao X, Ren S, Li J, Chen H, Bei W. 2016. Formate-tetrahydrofolate ligase is involved in the virulence of Streptococcus suis serotype 2. Microb Pathog 98:149–54.

71. Blanchette KA, Shenoy AT, Milner J, 2nd, Gilley RP, McClure E, Hinojosa CA, Kumar N, Daugherty SC, Tallon LJ, Ott S, King SJ, Ferreira DM, Gordon SB, Tettelin H, Orihuela CJ. 2016. Neuraminidase A-exposed galactose promotes Streptococcus pneumoniae biofilm formation during colonization. Infect Immun 84:2922–32.

72. Chimalapati S, Cohen JM, Camberlein E, MacDonald N, Durmort C, Vernet T, Hermans PW, Mitchell T, Brown JS. 2012. Effects of deletion of the Streptococcus pneumoniae lipoprotein diacylglyceryl transferase gene lgt on ABC transporter function and on growth in vivo. PLoS One 7:e41393.

73. Martin B, Prudhomme M, Alloing G, Granadel C, Claverys JP. 2000. Cross-regulation of competence pheromone production and export in the early control of transformation in Streptococcus pneumoniae. Mol Microbiol 38:867–78.

74. Granok AB, Parsonage D, Ross RP, Caparon MG. 2000. The RofA binding site in Streptococcus pyogenes is utilized in multiple transcriptional pathways. J Bacteriol 182:1529–40.

75. Havarstein LS, Coomaraswamy G, Morrison DA. 1995. An unmodified heptadecapeptide pheromone induces competence for genetic transformation in Streptococcus pneumoniae. Proc Natl Acad Sci U S A 92:11140–4.

76. Khandavilli S, Homer KA, Yuste J, Basavanna S, Mitchell T, Brown JS. 2008. Maturation of Streptococcus pneumoniae lipoproteins by a type II signal peptidase is required for ABC transporter function and full virulence. Mol Microbiol 67:541–57.

77. Domenech M, Ramos-Sevillano E, Garcia E, Moscoso M, Yuste J. 2013. Biofilm formation avoids complement immunity and phagocytosis of Streptococcus pneumoniae. Infect Immun 81:2606–15.

78. Sletvold H, Johnsen PJ, Wikmark OG, Simonsen GS, Sundsfjord A, Nielsen KM. 2010. Tn1546 is part of a larger plasmid-encoded genetic unit horizontally disseminated among clonal Enterococcus faecium lineages. Journal of Antimicrobial Chemotherapy 65:1894–1906.

79. Ewels P, Magnusson M, Lundin S, Kaller M. 2016. MultiQC: summarize analysis results for multiple tools and samples in a single report. Bioinformatics 32:3047–8.

80. Bolger AM, Lohse M, Usadel B. 2014. Trimmomatic: a flexible trimmer for Illumina sequence data. Bioinformatics 30:2114–20.

81. Langmead B, Salzberg SL. 2012. Fast gapped-read alignment with Bowtie 2. Nat Methods 9:357–9.

82. Li H, Handsaker B, Wysoker A, Fennell T, Ruan J, Homer N, Marth G, Abecasis G, Durbin R, Genome Project Data Processing S. 2009. The sequence alignment/map format and SAMtools. Bioinformatics 25:2078–9.

83. Nicol JW, Helt GA, Blanchard SG, Jr., Raja A, Loraine AE. 2009. The integrated genome browser: free software for distribution and exploration of genome-scale datasets. Bioinformatics 25:2730–1.

84. Liao Y, Smyth GK, Shi W. 2014. featureCounts: an efficient general purpose program for assigning sequence reads to genomic features. Bioinformatics 30:923–30.

85. Love MI, Huber W, Anders S. 2014. Moderated estimation of fold change and dispersion for RNA-seq data with DESeq2. Genome Biol 15:550.

86. Yu G, Wang LG, Han Y, He QY. 2012. clusterProfiler: an R package for comparing biological themes among gene clusters. OMICS 16:284–7.

