## Supplementary figures and images for "Essential role of proline synthesis and the one-carbon metabolism pathways for systemic virulence of *Streptococcus pneumoniae*"

### Supp Figures

**Supplementary Figure 1**

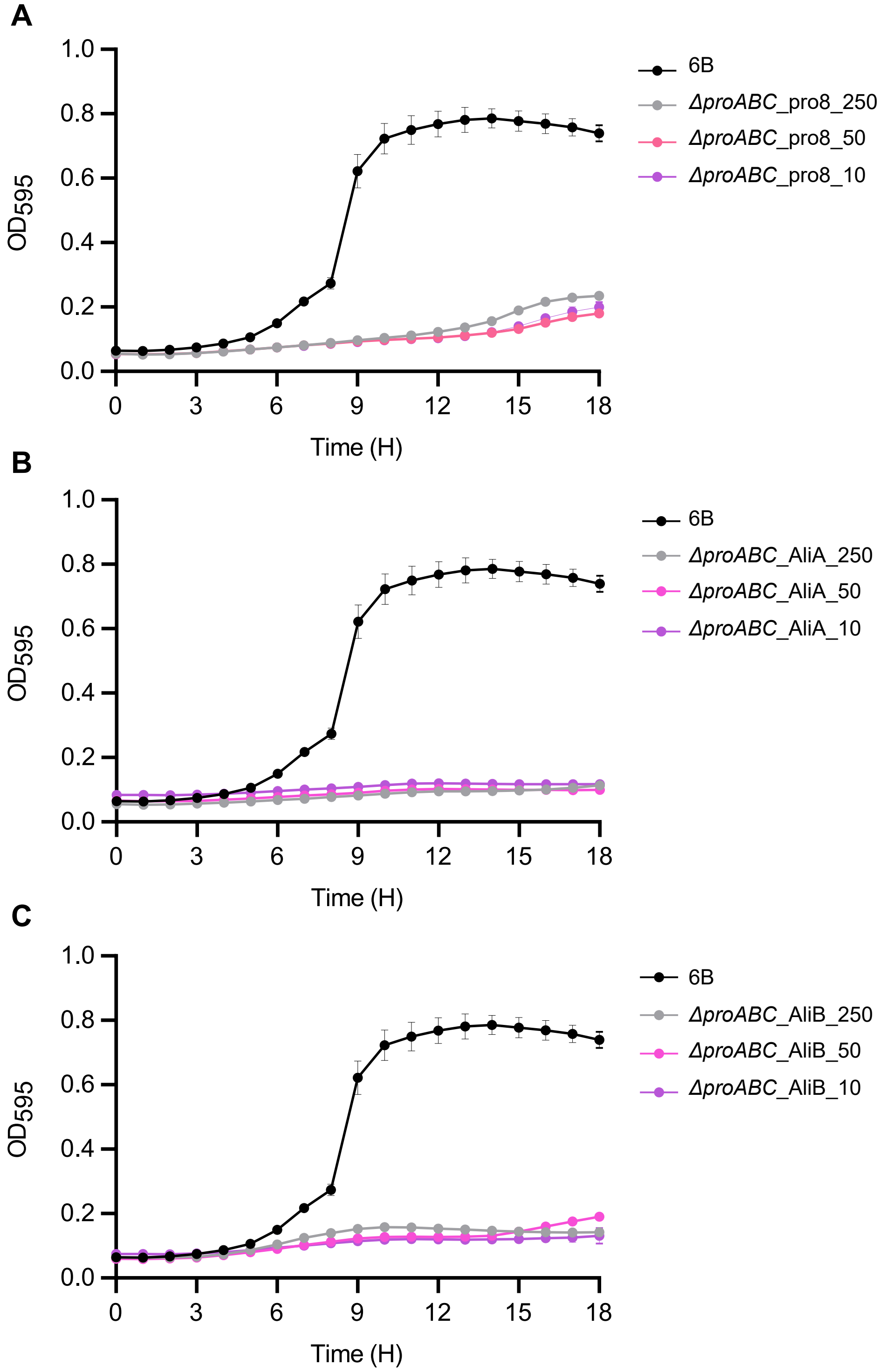

**Supplementary Figure 2**

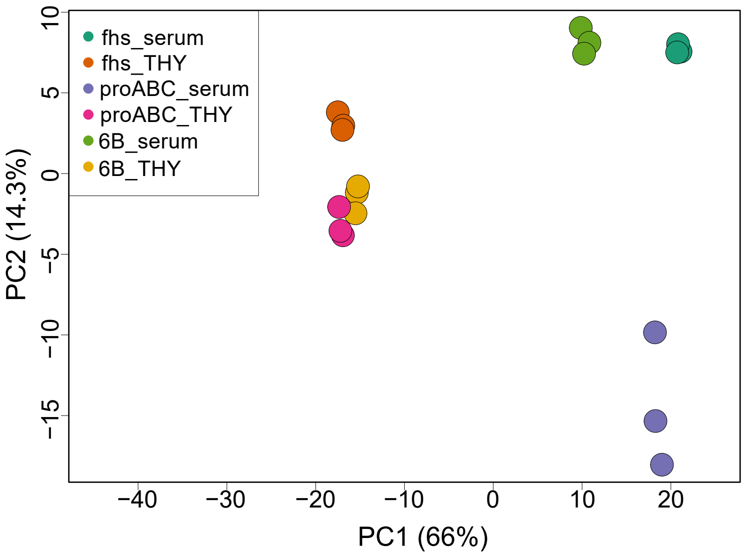
