## Supplementary material for "Essential role of proline synthesis and the one-carbon metabolism pathways for systemic virulence of *Streptococcus pneumoniae*": Supp Tables

**Supplementary tables**

**Supplementary Table 1.** Strains, primers and plasmids used in this study.

| **Strain** | Description |  |
| --- | --- | --- |
| BHN418 | *S. pneumoniae* Serotype 6B isolate |  |
| D39 | *S. pneumoniae* Serotype 2 isolate |  |
|  | Primer name | Primer sequence |
| fhs | Fhs_UpF  Fhs_UpspecF  Fhs_UpspecR  Fhs_DownspecF  Fhs_Downspec R  Fhs_DownR | ggcaagtgggtaattcttga  ctttcaaatctaacatatctctgatcccccgtttgattttt  aaaaatcaaacgggggatcagagatatgttagatttgaaag  ttggatccattccgcgtccgcttatttttgtgtacaatagt  actattgtacacaaaaataagcggacgcggaatggatccaa  caaggaggagttctgcaattt |
| spr832 | Spr832_UpF  Spr832_UpspecF  Spr832_UpspecR  Spr832_DownspecF  Spr832_Downspec R  Spr832_DownR | ccaaacgggtatcttgttacag  tgttattcatgttataatggagatcccccgtttgattt  atccattaaaaatcaaacgggggatctccattataacatgaat  aaaaattggatccattccgcgtcagctttgactgcctctttt  aaagaggcagtcaaagctgacgcggaatggatccaat  ggaaactaccaatgctgtcttgttt |
| cps | Cps_UpF  Cps_UpKanF  Cps_UpKanR  Cps_DownKanF  Cps_DownKanR  Cps_DownR | ggattgataaaggtattggtggt  gctttctgtgtggaattactataaatattgtcgatactatgttatacgccaac  gttggcgtataacatagtatcgacaatatttatagtaattccacacaga  cttttctgaagtacatccgcaacgaaaatgatgaaaagttcaaaac  gttttgaacttttcatcattttcgttgcggatgtacttcagaaaag  cagtttgtccattcaactgag |
| **Plasmids** | Description |  |
| pR412 | Derived from ColE1, carrying a 1145bp minitransposon that contains Himar1 IRs flanking *add9* gene. SpcR (1) |  |
| pPEPY | Complementation construct with a kan selectable marker. Descried by Jan-Willem Veening (Addgene plasmid #122633). It is a promoterless integrative plasmid (2) |  |
| pABG5mini | Derived from pMGC66, an *E. coli* streptococcal shuttle vector that harbors phoZF (3) |  |

**Supplementary Table 2.** Genes showing differential expression (log_2_ fold change) in the wild type 6B strain when cultured to mid-log growth phase in THY broth compared to human serum. Genes in bold are deleted in the mutant strains.

| **BHN418 gene number and name** | **TIGR4 gene number**  **(if known)** | **6B log_2_ fold change in serum v. THY** |
| --- | --- | --- |
| Spn_00006 | SP0508 | 2.074 |
| Spn_00015 | SP0498 | 1.875 |
| Spn_00023_hsdS_3 | SP0508 | 2.200 |
| Spn_00037_hsdS_6 |  | 2.271 |
| Spn_00041 |  | -2.513 |
| Spn_00042_rpsP_1 |  | -1.963 |
| Spn_00043_trxA_1 |  | -2.311 |
| Spn_00044_queF |  | -1.871 |
| Spn_00057 |  | -3.711 |
| Spn_00058 |  | -3.460 |
| Spn_00069 |  | 2.104 |
| Spn_00070 | SP1802 | 3.417 |
| Spn_00071 |  | 2.802 |
| Spn_00072 |  | 2.293 |
| Spn_00113 |  | 1.753 |
| Spn_00115_xpt | SP1847 | -2.832 |
| Spn_00116_ygfU | SP1848 | -3.485 |
| Spn_00124_xylB |  | -2.207 |
| Spn_00125_adhR |  | -2.038 |
| Spn_00126_czcD |  | -3.495 |
| Spn_00128_pnuC |  | -6.568 |
| Spn_00136_feuB | SP1869 | 2.553 |
| Spn_00137_fepD_1 | SP1870 | 2.483 |
| Spn_00138_fhuC | SP1871 | 2.476 |
| Spn_00139_yclQ | SP1872 | 2.566 |
| Spn_00150_treC | SP1883 | 4.551 |
| Spn_00151_treP | SP1884 | 4.498 |
| Spn_00152_treR | SP1885 | 2.205 |
| Spn_00160_amiA | SP1891 | -1.728 |
| Spn_00167_rhaS | SP1899 | -1.710 |
| Spn_00198_groEL | SP1906 | 2.116 |
| Spn_00207 | SP1917 | -1.914 |
| Spn_00208 | SP1917 | -1.571 |
| Spn_00227_dinF | SP1939 | 1.601 |
| Spn_00240_rpoC | SP1960 | 2.405 |
| Spn_00241_rpoB | SP1961 | 2.302 |
| Spn_00250_kdtB | SP1968 | 1.852 |
| Spn_00260_cbf1 | SP1980 | 1.858 |
| Spn_00261 | SP1981 | 1.620 |
| Spn_00268 | SP1988 | -1.676 |
| Spn_00269 | SP0109 | -2.656 |
| Spn_00286_ybhF | SP2003 | 1.874 |
| Spn_00306_pbp2a | SP2010 | 1.816 |
| Spn_00311_nadC | SP2016 | -4.442 |
| Spn_00312 | SP2017 | -2.727 |
| Spn_00315_gmuD_1 | SP2021 | -2.393 |
| Spn_00316_gmuC_3 | SP2022 | -1.923 |
| Spn_00317_licB_2 | SP2023 | -2.036 |
| Spn_00318_gmuA | SP2024 | -1.912 |
| Spn_00348 | SP1582 | 1.589 |
| Spn_00349_adh | SP2055 | -1.676 |
| Spn_00350_nagA | SP2056 | -1.837 |
| Spn_00351_oatA | SP2057 | 2.326 |
| Spn_00354_pcp_1 | SP2060 | -1.670 |
| Spn_00356_marR_1 | SP2062 | -1.961 |
| Spn_00358 | SP2063 | -2.610 |
| Spn_00393_pstS_1 | SP2084 | -2.933 |
| Spn_00413_glgP-2 | SP2106 | 1.893 |
| Spn_00414_malM | SP2107 | 1.616 |
| Spn_00425 | SP2116 | -2.128 |
| Spn_00426 | SP2117 | -2.560 |
| Spn_00427 | SP2118 | -3.142 |
| Spn_00428 | SP2119 | -3.392 |
| Spn_00429 | SP2120 | -3.064 |
| Spn_00434 | SP2125 | -2.788 |
| Spn_00435 | SP0174 | -2.805 |
| Spn_00436_ilvD | SP2126 | -4.010 |
| Spn_00442 | PS2132 | -3.517 |
| Spn_00443 | SP2133 | -2.766 |
| Spn_00444_rpmF | SP2134 | -1.782 |
| Spn_00445_rpmGA | SP2135 | -1.510 |
| Spn_00446_pcpA | SP2136 | 2.025 |
| Spn_00448 | SP2137 | 1.931 |
| Spn_00451 | SP2141 | 2.227 |
| Spn_00452_bglK_1 | SP2142 | 1.534 |
| Spn_00453_mngB | SP2143 | 2.653 |
| Spn_00454 | SP2144 | 2.007 |
| Spn_00482_dltD | SP2173 | 3.183 |
| Spn_00483_dltC | SP2174 | 2.821 |
| Spn_00484_dltB | SP2175 | 2.950 |
| Spn_00485_dltA | SP2176 | 2.763 |
| Spn_00499_bag | SP2190 | -1.808 |
| Spn_00506_yycG | SP2192 | 1.772 |
| Spn_00527 | SP1442 | -2.105 |
| Spn_00528_tsf | SP2214 | -2.390 |
| Spn_00531_mreD | SP2217 | 2.758 |
| Spn_00532_mreC | SP2218 | 2.495 |
| Spn_00533_ecfT | SP2219 | 1.985 |
| Spn_00534_ecfA2 | SP2220 | 2.018 |
| Spn_00535_ecfA1 | SP2221 | 2.053 |
| Spn_00536_pgsA | SP2222 | 2.150 |
| Spn_00537 | SP2223 | 2.117 |
| Spn_00538 | SP2224 | 2.738 |
| Spn_00539 | SP2225 | 2.178 |
| Spn_00544_yheS_1 | SP2230 | 1.682 |
| Spn_00545 | SP2231 | 3.596 |
| Spn_00555_htrA | SP2239 | 1.515 |
| Spn_00556_parB | SP2240 | 1.759 |
| Spn_00561_pth | SP0005 | 1.723 |
| Spn_00562_trcF | SP0006 | 3.150 |
| Spn_00566 | SP0010 | 1.849 |
| Spn_00567_tilS | SP0011 | 2.407 |
| Spn_00568_hgt | SP0012 | 2.261 |
| Spn_00569_ftsH | SP0013 | 2.363 |
| Spn_00580_tadA | SP0020 | 2.054 |
| Spn_00591_polA | SP0032 | 1.578 |
| Spn_00593_yeiH | SP0034 | 1.931 |
| Spn_00601_comA_1 | SP0042 | -1.988 |
| Spn_00602_comB | SP0043 | -1.797 |
| Spn_00603_purC | SP0044 | -2.308 |
| Spn_00604_purL | SP0045 | -3.442 |
| Spn_00605_purF | SP0046 | -2.679 |
| Spn_00606_purM | SP0047 | -2.706 |
| Spn_00607_purN | SP0048 | -2.316 |
| Spn_00609_purH | SP0050 | -2.256 |
| Spn_00611_purE | SP0053 | -1.738 |
| Spn_00613 | SP0055 | 1.815 |
| Spn_00616_frlR | SP0058 | -1.664 |
| Spn_00617_bgaC | SP0060 | 2.837 |
| Spn_00618_PTS-EIIB | SP0061 | 2.073 |
| Spn_00619_PTS-EIIC | SP0062 | 1.588 |
| Spn_00620_manZ_2 | SP0063 | 1.763 |
| Spn_00621_PTS-EII_2 | SP0064 | 1.515 |
| Spn_00622_agaS | SP0265 | 2.117 |
| Spn_00623_mro | SP0266 | 2.071 |
| Spn_00638_rpsD | SP0085 | -2.989 |
| Spn_00640_ugpA_1 | SP0090 | 2.350 |
| Spn_00641_ycjP_2 | SP0091 | 2.419 |
| Spn_00642 | SP0092 | 2.190 |
| Spn_00645 | SP0097 | -1.537 |
| Spn_00650_pgaC | SP0102 | 2.179 |
| Spn_00651_capD | SP0103 | 2.048 |
| Spn_00655_xlyA | SP0107 | -2.288 |
| Spn_00656 | SP0109 | -1.585 |
| Spn_00659_artP_1 | SP0112 | -3.108 |
| Spn_00660_argG | SP0113 | -3.198 |
| Spn_00676_mnmA | SP0118 | -2.435 |
| Spn_00698_mutR | SP0141 | -4.152 |
| Spn_00699 | SP0142 | -4.267 |
| Spn_00700 | SP0143 | -2.876 |
| Spn_00701 | SP0144 | -1.974 |
| Spn_00702 | SP0145 | -1.687 |
| Spn_00716 | SP0159 | -4.321 |
| Spn_00717 | SP0160 | -1.762 |
| Spn_00718 | SP0161 | -2.026 |
| Spn_00720 | SP0164 | -1.688 |
| Spn_00721_epiD | SP0165 | -2.098 |
| Spn_00738_Int-Tn | SP1129 | 2.273 |
| Spn_00739 Prophage element | SP0727 | 2.625 |
| Spn_00740 | _ | 2.438 |
| Spn_00755_pepP | SP0187 | 1.684 |
| Spn_00759_yrrK | _ | -1.730 |
| Spn_00760 | SP0194 | -1.502 |
| Spn_00766_nrdD_1 | SP0202 | -1.787 |
| Spn_00771_rplC | SP0209 | -1.642 |
| Spn_00772_rplD | SP0210 | -1.922 |
| Spn_00773_rplW | SP0211 | -1.654 |
| Spn_00774_rplB | SP0212 | -1.986 |
| Spn_00775_rpsS | SP0213 | -1.824 |
| Spn_00776_rplV | SP0214 | -1.641 |
| Spn_00777_rpsC | SP0215 | -1.935 |
| Spn_00778_rplP | SP0216 | -2.134 |
| Spn_00779_rpmC | SP0217 | -1.857 |
| Spn_00780_rpsQ | SP0218 | -2.035 |
| Spn_00781_rplN | SP0219 | -2.347 |
| Spn_00782_rplX | SP0220 | -2.093 |
| Spn_00783_rplE | SP0221 | -2.122 |
| Spn_00784_rpsN2 | SP0222 | -2.309 |
| Spn_00785_rpsH | SP0224 | -2.008 |
| Spn_00786_rplF | SP0225 | -2.139 |
| Spn_00787_rplR | SP0226 | -1.929 |
| Spn_00788_rpsE | SP0227 | -2.147 |
| Spn_00789_rpmD | SP0228 | -1.723 |
| Spn_00790_rplO | SP0229 | -2.470 |
| Spn_00791_secY_1 | SP0230 | -2.079 |
| Spn_00792_adk | SP0231 | 1.705 |
| Spn_00813_deoR | SP0246 | -1.651 |
| Spn_00838_polC_1 | SP0274 | 1.669 |
| Spn_00839 | SP0275 | -3.527 |
| Spn_00840_yafQ | SP0276 | -3.107 |
| Spn_00841_polC_2 | SP277 | -2.770 |
| Spn_00842_pepS | SP0278 | -3.166 |
| Spn_00843 | SP0279 | -2.532 |
| Spn_00844_rsuA1 | SP0280 | -1.637 |
| Spn_00845_pepC | SP0281 | 2.296 |
| Spn_00849_adhP | SP0285 | -1.865 |
| Spn_00850_yidA_1 | SP0286 | 2.806 |
| Spn_00851_pbuO | SP0287 | -2.443 |
| Spn_00859_rplM | SP0294 | -1.738 |
| Spn_00860_rpsI | SP0295 | -2.004 |
| Spn_00862 | SP0878 | -1.748 |
| Spn_00863 | _ | -3.138 |
| Spn_00864 | _ | -3.582 |
| Spn_00870_yicI | SP0312 | 2.174 |
| Spn_00887 | SP2241 | -2.288 |
| Spn_00888 | SP0332 | -2.337 |
| Spn_00889 | SP0332 | -2.330 |
| Spn_00890 | SP0333 | -1.624 |
| Spn_00914_aliA | SP0366 | -5.164 |
| Spn_00915 | SP0368 | 1.682 |
| Spn_00916 | SP1262 | 2.615 |
| Spn_00920_gpsB | SP0372 | -1.782 |
| Spn_00932_mvd1 | SP0382 | 1.781 |
| Spn_00933 | SP0383 | 2.319 |
| Spn_00934_fni | SP0384 | 2.192 |
| Spn_00935 | SP0385 | 3.255 |
| Spn_00936_vraS | SP0386 | 3.204 |
| Spn_00937_vraR_2 | SP0387 | 3.043 |
| Spn_00938 | SP0389 | 2.418 |
| Spn_00939_cbpG | SP0390 | 3.004 |
| Spn_00940_lytA_3 | SP0391 | 1.910 |
| Spn_00944_mtlA | SP0394 | -2.639 |
| Spn_00945 | SP0395 | -1.940 |
| Spn_00946_mtlF | SP0396 | -1.596 |
| Spn_00947_mtlD | SP0397 | -1.768 |
| Spn_00963_fabM | SP0415 | 2.326 |
| Spn_00976 | SP1262 | -2.817 |
| Spn_00992_ilvB | SP0445 | -2.747 |
| Spn_00993_ilvH | SP0446 | -2.230 |
| Spn_00994_ilvC | SP0447 | -2.312 |
| Spn_00995 | SP0450 | -1.923 |
| Spn_00997_ilvA | SP0450 | -2.727 |
| Spn_01007_pfl | SP0459 | 2.023 |
| Spn_01008 | SP0460 | 1.876 |
| Spn_01021 | SP0471 | -1.695 |
| Spn_01024_trkG2 | SP0479 | 2.462 |
| Spn_01025_trkA2 | SP0480 | 2.511 |
| Spn_01035 | SP0489 | 1.771 |
| Spn_01036 | SP0490 | 1.827 |
| Spn_01037 | SP0490 | 2.068 |
| Spn_01066_hrcA_1 | SP0515 | 2.738 |
| Spn_01067_grpE_1 | SP0516 | 2.265 |
| Spn_01068_dnaK_1 | SP0517 | 2.792 |
| Spn_01069 | SP0518 | 3.219 |
| Spn_01070_dnaJ_1 | SP0519 | 2.960 |
| Spn_01073_ecsA_1 | SP0522 | 1.728 |
| Spn_01074 | SP0523 | 2.059 |
| Spn_01075 | SP0523 | 1.908 |
| Spn_01081_dnaK_2 | SP0517 | 3.337 |
| Spn_01082_dnaK_3 | SP0517 | -4.313 |
| Spn_01088 | SP0523 | 2.295 |
| Spn_01106_blpN_2 | SP0540 | -1.544 |
| Spn_01127 | SP0565 | 2.943 |
| Spn_01128_ydaF_1 | SP0566 | 2.629 |
| Spn_01129 | SP0567 | 3.096 |
| Spn_01130_valS | SP0568 | 2.865 |
| Spn_01139_pheS | SP0579 | 2.565 |
| Spn_01140_paiA | SP0580 | 2.381 |
| Spn_01141_pheT_2 | SP0581 | 3.185 |
| Spn_01158_regX3 | SP0601 | 1.600 |
| Spn_01159_vncS | SP0604 | 1.920 |
| Spn_01161_ndoR | SP0606 | -1.817 |
| Spn_01162_yecS_2 | SP0607 | -2.262 |
| Spn_01163_artQ_1 | SP0608 | -2.595 |
| Spn_01164_peb1A_1 | SP0609 | -2.220 |
| Spn_01165_peb1A_2 | SP0609 | -1.603 |
| Spn_01166_glnQ_1 | SP0610 | -1.721 |
| Spn_01171_fibB | SP0616 | 2.171 |
| Spn_01172 | SP0617 | 2.354 |
| Spn_01180_brnQ | SP0626 | 2.970 |
| Spn_01193 | SP0645 | 2.293 |
| Spn_01194 | SP0646 | 1.523 |
| Spn_01205_nhaK | SP0654 | 1.669 |
| Spn_01206 | SP0657 | 1.816 |
| Spn_01213_zmpB | SP0664 | 1.587 |
| Spn_01221_hflX | SP0672 | 1.698 |
| Spn_01222 | SP0672 | 2.268 |
| Spn_01223_rnz | SP0674 | 2.008 |
| Spn_01224 | SP0675 | 2.337 |
| Spn_01229_rsuA | SP0680 | -2.200 |
| Spn_01231 | SP0682 | 2.162 |
| Spn_01240_pyrF | SP0701 | -2.880 |
| Spn_01241_pyrE | SP0702 | -2.631 |
| Spn_01269_copY | SP0727 | -1.829 |
| Spn_01270 | SP0728 | -1.695 |
| Spn_01272_spxB | SP0730 | 2.554 |
| Spn_01273 | SP0731 | 2.072 |
| Spn_01289_gmuF | SP0736 | -1.544 |
| Spn_01295 | SP0742 | 1.774 |
| Spn_01301_livJ | SP0749 | -3.383 |
| Spn_01302_livH | SP0750 | -2.257 |
| Spn_01303_livM | SP0751 | -2.032 |
| Spn_01304_lptB | SP0752 | -1.866 |
| Spn_01305_livF | SP0753 | -2.185 |
| Spn_01319 | SP0769 | 1.656 |
| Spn_01320 | SP0770 | 1.652 |
| Spn_01333_gor | SP0784 | 2.081 |
| Spn_01334 | SP0785 | -2.569 |
| Spn_01335_salX | SP0786 | -2.332 |
| Spn_01336_macB_4 | _ | -2.067 |
| Spn_01340_ydhF | SP0791 | 1.865 |
| Spn_01346_pepN | SP0797 | 1.705 |
| Spn_01349 | SP0800 | -2.580 |
| Spn_01352_rodA | SP0803 | 1.614 |
| Spn_01366_clpE | SP0717 | 1.796 |
| Spn_01372_rpiA | SP0828 | 1.743 |
| Spn_01373_deoB | SP0724 | 1.591 |
| Spn_01374 | SP0830 | 1.748 |
| Spn_01376_flaR | SO0832 | 1.533 |
| Spn_01378_deoD_2 | SP0837 | 1.699 |
| Spn_01398_parC | SP0855 | 1.758 |
| Spn_01399_ilvE | SP0856 | -2.123 |
| Spn_01400_sarA_1 | SP0857 | -1.886 |
| Spn_01401 | SP0858 | -2.377 |
| Spn_01402 | SP0859 | -1.952 |
| Spn_01403_pcp_2 | SP0860 | -1.839 |
| Spn_01412_dnaX | SP0865 | 2.221 |
| Spn_01413 | SP0866 | 2.727 |
| Spn_01416_csd | SP0869 | 1.682 |
| Spn_01417_nifU | SP0870 | 2.693 |
| Spn_01418 | SP0871 | 2.812 |
| Spn_01419_pbp3 | SP0872 | 1.760 |
| Spn_01423_fruR | SP0875 | 5.280 |
| Spn_01424_fruB | SP0876 | 4.987 |
| Spn_01425_fruA_1 | SP0877 | 4.690 |
| Spn_01446_pyk | SP0897 | 1.962 |
| Spn_01447 | SP0899 | -2.655 |
| Spn_01449 | SP0897 | -2.203 |
| Spn_01450 | SP0899 | -2.518 |
| Spn_01451 | SP0897 | 2.335 |
| Spn_01452 | SP0899 | 2.034 |
| Spn_01498_rpmI | SP0960 | -1.730 |
| Spn_01499_rplT | SP0961 | -2.304 |
| Spn_01501_pyrDII | SP0963 | -4.456 |
| Spn_01502_pyrD | SP0964 | -3.862 |
| Spn_01503_lytB_3 | SP0965 | -2.772 |
| Spn_01508_mutM | SP0970 | 2.120 |
| Spn_01509_coaE | SP0971 | 1.663 |
| Spn_01510_tetA | SP0972 | 2.454 |
| Spn_01530_ugpQ | SP0994 | 2.029 |
| Spn_01533_dsbD_2 | SP0999 | 2.566 |
| Spn_01534 | SP1000 | 2.851 |
| Spn_01552_tdk | SP1018 | -1.729 |
| Spn_01553_prfA | SP1020 | -1.687 |
| Spn_01555_rimN | SP1022 | -1.577 |
| Spn_01562 |  | -2.336 |
| Spn_01564_fepD_2 |  | 2.007 |
| Spn_01565_yfhA |  | 2.455 |
| Spn_01566_yusV |  | 2.010 |
| Spn_01573 | SP1042 | -3.132 |
| Spn_01574 | SP1043 | -4.736 |
| Spn_01575 | SP1044 | -4.631 |
| Spn_01576 | SP1045 | -2.968 |
| Spn_01597_obg | SP1079 | -1.754 |
| Spn_01608_rex | SP1090 | -1.993 |
| Spn_01615_ppnK1 | SP1098 | 1.721 |
| Spn_01616_rluA3 | SP1099 | 2.054 |
| Spn_01617_pta | SP1100 | 1.602 |
| Spn_01620_rplU | SP1105 | -1.926 |
| Spn_01621 | SP1106 | -2.108 |
| Spn_01622_rpmA | SP1107 | -2.027 |
| Spn_01630_ligA | SP1117 | 2.301 |
| Spn_01631_pulA_2 | SP1118 | 3.040 |
| Spn_01632_gapN | SP1119 | -4.014 |
| Spn_01639 | SP1127 | 2.091 |
| Spn_01640_eno | SP1128 | 1.709 |
| Spn_01643 | SP0299 | -1.603 |
| Spn_01645_rexA | SP1152 | 1.555 |
| Spn_01646 | SP1153 | 1.513 |
| Spn_01647_iga_2 | SP1154 | 3.854 |
| Spn_01656 | _ | -2.457 |
| Spn_01657 | SP1589 | -2.902 |
| Spn_01658_traG | _ | -2.691 |
| Spn_01659 | SP0668 | -2.820 |
| Spn_01660 | SP2182 | -2.605 |
| Spn_01662 | SP1334 | -2.371 |
| Spn_01663 | SP2216 | -1.693 |
| Spn_01672 |  | -2.685 |
| Spn_01673 |  | -1.848 |
| Spn_01674 |  | -1.844 |
| Spn_01675 |  | -1.579 |
| Spn_01676 |  | -1.934 |
| Spn_01677_macB_6 | SP0912 | -2.885 |
| Spn_01678 | SP0913 | -1.979 |
| Spn_01687 | SP1058 | -2.008 |
| Spn_01688 | SP1059 | -2.151 |
| Spn_01699_xerS | SP1159 | -3.475 |
| Spn_01740_recN | SP1202 | 1.631 |
| Spn_01741_ahrC | SP1203 | 1.555 |
| Spn_01756_ldhA | SP1220 | 1.684 |
| Spn_01758 | SP1223 | -1.797 |
| Spn_01759 | SP1224 | -2.044 |
| Spn_01774 |  | -1.728 |
| Spn_01775_glnP |  | -1.866 |
| Spn_01776_artM_4 |  | -1.865 |
| Spn_01778_ftsY |  | 1.950 |
| Spn_01779_yidA_3 |  | 1.817 |
| Spn_01780_ybjI | SP1247 | 2.483 |
| Spn_01781_smc | SP1248 | 2.055 |
| Spn_01784_guaC | SP1249 | -1.937 |
| Spn_01797 | sp1264 | -1.608 |
| Spn_01804_tacF |  | 1.648 |
| Spn_01805_licD1 | SP1273 | 2.324 |
| Spn_01806_licD2 | SP1274 | 2.088 |
| Spn_01807_carB | SP1275 | -3.503 |
| Spn_01808_carA | SP1276 | -3.818 |
| Spn_01809_pyrB | SP1277 | -3.629 |
| Spn_01810_pyrR | SP1278 | -3.397 |
| Spn_01815_yheS_2 | SP1282 | -1.879 |
| Spn_01819_pyrP | SP1286 | -4.023 |
| Spn_01825 |  | 1.604 |
| Spn_01838_gdhA | SP1306 | -2.178 |
| Spn_01866 | SP1363 | 2.488 |
| Spn_01867 | SP1364 | 1.962 |
| Spn_01876_tyrC | sp1373 | 1.676 |
| Spn_01882 | SP1380 | 1.683 |
| Spn_01885_alaS | SP1384 | 2.573 |
| Spn_01886 |  | 2.483 |
| Spn_01895_yjiR | SP1393 | -2.749 |
| Spn_01896_glnH | SP1394 | -3.617 |
| Spn_01916_rpsU |  | -2.125 |
| Spn_01917_nagB | SP1415 | -2.610 |
| Spn_01919_lytA_5 | SP1416 | -1.645 |
| Spn_01921_ydaF_4 | SP1419 | -2.013 |
| Spn_01922_nadE |  | -3.332 |
| Spn_01923_pncB2 |  | -3.329 |
| Spn_01924 | SP1422 | -3.186 |
| Spn_01925 |  | -3.930 |
| Spn_01926 | SP1424 | -3.362 |
| Spn_01927_artM_5 |  | -3.685 |
| Spn_01929 |  | -2.070 |
| Spn_01930_yhbU_2 |  | -3.547 |
| Spn_01936_yicL_2 |  | 1.887 |
| Spn_01937 |  | 1.633 |
| Spn_01954_pdxT |  | -4.318 |
| Spn_01955_pdx1 | SP1468 | -4.906 |
| Spn_01960_ynzC | SP1473 | 2.278 |
| Spn_01961_glyS | SP1474 | 2.751 |
| Spn_01962_glyQ | SP1475 | 2.521 |
| Spn_01963 | SP1476 | 2.482 |
| Spn_01964 | SP1477 | 2.085 |
| Spn_01980_artP_2 |  | -1.647 |
| Spn_01981_glnQ_5 |  | -2.033 |
| Spn_01982_yecS_3 |  | -1.883 |
| Spn_01983 |  | 2.657 |
| Spn_01984_yhhT_2 |  | 2.388 |
| Spn_01986_atpC | SP1507 | 2.341 |
| Spn_01987_atpD | SP1508 | 2.170 |
| Spn_01988_atpG | SP1509 | 2.588 |
| Spn_01989_atpA | Sp1510 | 2.379 |
| Spn_01990_atpH | sp1511 | 2.209 |
| Spn_01991_atpF | sp1512 | 1.848 |
| Spn_01992_atpB | SP1513 | 2.272 |
| Spn_01993_atpE | Sp1514 | 1.808 |
| Spn_01994 |  | 1.758 |
| Spn_01995 |  | 1.546 |
| Spn_02005_lmrA_2 | too short | -1.636 |
| Spn_02006_sarA_2 | SP1527 | -3.866 |
| Spn_02025 |  | -1.982 |
| Spn_02028 |  | -2.266 |
| Spn_02029 |  | -1.585 |
| Spn_02047_engB |  | 1.924 |
| Spn_02048_clpX | SP1569 | 1.865 |
| Spn_02049 |  | 1.590 |
| Spn_02051_dps |  | -2.511 |
| Spn_02054_dnaD |  | 2.077 |
| Spn_02061_codY | SP1584 | -1.666 |
| Spn_02064_lpd |  | 3.114 |
| Spn_02067_pepQ | SP1591 | 2.053 |
| Spn_02072_hmpT |  | -2.681 |
| Spn_02073_pdxK |  | -3.084 |
| Spn_02074_truA | SP1599 | -2.490 |
| Spn_02077 |  | -1.554 |
| Spn_02087 | SP1612 | -2.369 |
| Spn_02088 | SP1612 | -1.516 |
| Spn_02104 |  | 2.266 |
| Spn_02105_thrS | SP1631 | 2.491 |
| Spn_02115_relA | SP1645 | -1.985 |
| Spn_02117_pepO | SP1647 | 2.014 |
| Spn_02118 | SP1648? | -2.573 |
| Spn_02119_psaC | SP1649 | -2.304 |
| Spn_02120_psaA | SP1650 | -1.826 |
| Spn_02123_macB_7 | SP1653 | -2.235 |
| Spn_02128 | SP1658 | -1.815 |
| Spn_02130_ileRS | SP1659 | 4.242 |
| Spn_02132_divIVA |  | 1.995 |
| Spn_02133 |  | 2.104 |
| Spn_02139 |  | 1.508 |
| Spn_02143_recR | SP1672 | 1.552 |
| Spn_02154_yesO_1 | SP1683 | -1.754 |
| Spn_02156_nanE2 | SP1685 | -1.772 |
| Spn_02164_nanA_5 | SP1692-93 | -1.870 |
| Spn_02199_mvaA | SP1726 | 2.090 |
| Spn_02207_rsmB | SP1734 | -1.870 |
| Spn_02208_fmt | SP1735 | -2.299 |
| Spn_02209_priA | SP0828 | -2.729 |
| Spn_02210_rpoZ | SP1737 | -1.620 |
| Spn_02215 |  | 1.818 |
| Spn_02216 |  | 1.758 |
| Spn_02218_yecD_2 |  | 1.609 |
| Spn_02226_sstT |  | -2.275 |
| Spn_02231 | SP1757 | 1.811 |
| Spn_02232_tagE | SP1758 | 1.881 |
| Spn_02233_secA2 | SP1759 | 1.993 |
| Spn_02235_asp2 | SP1761 | 2.189 |
| Spn_02236 | SP1762 | 1.976 |
| Spn_02237_secY_2 | SP1763 | 1.903 |
| Spn_02238 | SP1762 | 1.605 |
| Spn_02239_gspA_2 | SP1761 | 2.096 |
| Spn_02240_gspA_3 | SP1760 | 2.114 |

**Supplementary Table 3.** Genes showing differential expression (log_2_ fold change) between the mutant strains and the wild type 6B strain when cultured to mid-log growth phase in THY broth. Genes in bold are deleted in the mutant strains (*proABC,* Spn_01479-81; *fhs*, Spn_01764).

| **BHN418 gene number and name** | **TIGR4 gene number**  **(if known)** | ***∆fhs* log_2_ fold change v. wild type** | ***∆proABC* log_2_ fold change v. wild type** |
| --- | --- | --- | --- |
| Spn_00006 | SP0508 |  | 1.665 |
| Spn_00012 | SP0500 | -1.951 |  |
| Spn_00015 | SP498 | -2.515 |  |
| Spn_00016 | SP498 | -2.041 |  |
| Spn_00136_feuB | SP1869 | 1.785 |  |
| Spn_00137_fepD_1 | SP1870 | 1.811 |  |
| Spn_00138_fhuC | SP1871 | 1.773 |  |
| Spn_00139_yclQ | SP1872 | 1.793 |  |
| Spn_00153 |  | 1.839 |  |
| Spn_00198_groEL | SP1906 | -2.853 | -1.781 |
| Spn_00199_groES | SP1907 | -2.760 | -1.955 |
| Spn_00256_misCB | SP1975 | -1.688 | -1.843 |
| Spn_00311_nadC | SP2016 | 1.812 |  |
| Spn_00319_adhE | SP2026 | -2.024 |  |
| Spn_00358 | SP2063 | -1.888 | -1.613 |
| Spn_00394_pstC_1 | SP2085 |  | 1.7222 |
| Spn_00395_pstA_1 | SP2086 |  | 1.823 |
| Spn_00396_pstB3_1 | SP2087 |  | 1.783 |
| Spn_00397_phoU_1 | SP2088 |  | 1.850 |
| Spn_00413_glgP-2 | SP2106 | -1.596 |  |
| Spn_00415_malX | SP2108 |  | 1.541 |
| Spn_00416_malF | SP2109 |  | 1.640 |
| Spn_00417_malG | SP2110 |  | 1.917 |
| Spn_00434 | SP2125 | 2.846 |  |
| Spn_00451 | sp2141 | -1.661 |  |
| Spn_00457_arcA | SP2148 | 2.900 |  |
| Spn_00458_arcB | SP2150 | 2.821 |  |
| Spn_00459_arcC | SP2151 | 1.823 |  |
| Spn_00460 | SP2152 | 1.583 |  |
| Spn_00495_glpK | SP2186 |  | -1.682 |
| Spn_00520 | SP2206 | -1.762 | 3.741 |
| Spn_00546 | SP2233 | 2.857 |  |
| Spn_00555_htrA | SP2239 |  | -2.066 |
| Spn_00556_parB | SP2240 |  | -2.162 |
| Spn_00641_ycjP_2 | SP0091 | -1.958 |  |
| Spn_00642 | SP0092 | -2.571 |  |
| Spn_00655_xlyA | SP_0107 | -3.235 | -2.297 |
| Spn_00698_mutR | SP0141 | 1.988 |  |
| Spn_00699 | SP0142 | 2.065 |  |
| Spn_00700 | SP0143 | 2.126 |  |
| Spn_00701 | SP0144 | 2.252 |  |
| Spn_00702 | SP0145 | 2.267 |  |
| Spn_00716 | SP0159 | 3.994 | 1.950 |
| Spn_00757 | SP0191 | -1.501 | -1.544 |
| Spn_00770_rpsJ | SP0208 |  |  |
| Spn_00816_gmuB | SP0249 | 1.888 |  |
| Spn_00817_gmuC_4 | SP0250 | 2.483 |  |
| Spn_00818_hpdB | SP0251 | 2.330 |  |
| Spn_00819_fsaA | SP0252 | 1.981 |  |
| Spn_00820_gldA | SP0253 | 1.893 |  |
| Spn_00846_manZ_3 | SP0282 |  | 1.683 |
| Spn_00847_manY_2 | SP0283 |  | 1.701 |
| Spn_00848_manL | SP0284 |  | 1.643 |
| Spn_00895_clpC_2 | SP0338 | -3.676 | -3.033 |
| Spn_00914_aliA | SP0366 | 4.063 | 1.783 |
| Spn_00935 | SP0385 | -2.077 |  |
| Spn_00936_vraS | SP0386 | -2.072 |  |
| Spn_00937_vraR_2 | SP0387 | -2.029 |  |
| Spn_00938 | SP0389 | -1.955 |  |
| Spn_00939_cbpG | SP0390 | -1.928 |  |
| Spn_00940_lytA_3 | SP0391 | -1.762 |  |
| Spn_00963_fabM | SP0415 | 1.679 |  |
| Spn_00964_marR_2 | SP0416 | 1.903 | 2.490 |
| Spn_00965_fabH | SP0417 | 1.940 | 2.494 |
| Spn_00966_acpP_2 | SO0418 |  | 1.769 |
| Spn_00967 | SP0419 | 3.260 | 3.517 |
| Spn_00968_fabD | SP0420 | 3.269 | 3.522 |
| Spn_00969_fabG | SP0421 | 3.081 | 3.350 |
| Spn_00970_fabF | SP0422 | 2.903 | 3.211 |
| Spn_00971_accB | SP0423 | 2.832 | 3.253 |
| Spn_00972_fabZ | SP0424 | 2.879 | 3.236 |
| Spn_00973_accC | SP0425 | 2.907 | 3.283 |
| Spn_00974_accD | SP0426 | 2.927 | 3.258 |
| Spn_00975_accA | SP0427 | 2.859 | 3.231 |
| Spn_00976 | SP1262 | 3.822 | 4.131 |
| Spn_00992_ilvB | SP0445 | 3.575 |  |
| Spn_00993_ilvH | SP0446 | 3.221 |  |
| Spn_00994_ilvC | SP0447 | 3.107 |  |
| Spn_00995 | SP0448 | 2.798 |  |
| Spn_00996 | SP0449 | 2.221 |  |
| Spn_00997_ilvA | SP0450 | 3.046 |  |
| Spn_01007_pfl | SP0459 | -1.559 |  |
| Spn_01011 | SP0463 | -1.998 |  |
| Spn_01012 | SP0464 | -1.715 |  |
| Spn_01030_cbiQ | SP0483 | 2.636 | 2.9890 |
| Spn_01032_cspR | SP0486 | 1.659 |  |
| Spn_01052_nanA_3 | SP0498 | -2.253 |  |
| Spn_01066_hrcA_1 | SP0515 | -3.826 | -2.261 |
| Spn_01067_grpE_1 | SP0516 | -4.355 | -2.905 |
| Spn_01068_dnaK_1 | SP0517 | -4.726 | -2.931 |
| Spn_01069 | SP0518 | -4.817 | -2.551 |
| Spn_01070_dnaJ_1 | SP0519 | -3.965 | -2.153 |
| Spn_01172 | SP0617 |  | -1.846 |
| Spn_01081_dnaK_2 | SP0517 | -4.927 | -3.010 |
| Spn_01082_dnaK_3 | SP0517 |  | 3.599 |
| Spn_01195_gatC_2 | SP0438 | -1.941 |  |
| Spn_01197_lacZ | SP0648 | -1.827 |  |
| Spn_01295 | SP0742 | -2.441 | -2.285 |
| Spn_01310_ptsG_1 | SP0758 |  | 1.643 |
| Spn_01311 | SP0759/60 |  | 1.502 |
| Spn_01347_ciaR | SP0798 |  | -1.657 |
| Spn_01348_ciaH | SP0799 |  | -1.743 |
| Spn_01399_ilvE | SP0856 | 1.810 |  |
| Spn_01400_sarA_1 | SP0857 | 1.807 |  |
| Spn_01401 | SP0858 | 1.710 |  |
| Spn_01402 | SP0859 | 1.712 |  |
| Spn_01403_pcp_2 | SP0860 | 1.533 |  |
| Spn_01423_fruR | SP0875 |  | -3.481 |
| Spn_01424_fruB | SP0876 |  | -3.810 |
| Spn_01425_fruA_1 | SP0877 |  | -3.506 |
| Spn_01427 | SP0879 | -2.700 | -2.295 |
| Spn_01440 | SP0891 | -1.804 |  |
| Spn_01456 | SP0910 | -3.067 | -2.519 |
| Spn_01457 | SP0911 | -1.510 | -1.575 |
| Spn_01459_macB_5 | SP0912 | -2.249 | -1.585 |
| Spn_01460 | SP0913 | -2.239 | -1.668 |
| Spn_01464_speA | SP0916 |  | 1.789 |
| Spn_01466_speE | SP0918 |  | 1.723 |
| Spn_01467 | SP0919 |  | 1.592 |
| Spn_01468_lysA_3 | SP0920 |  | 1.731 |
| Spn_01469_aguA | SP0921 |  | 1.604 |
| Spn_01470 | SP0922 |  | 1.579 |
| **Spn_01479_proB** | SP0931 |  | -5.667 |
| **Spn_01480_proA** | SP0932 |  | -6.160 |
| **Spn_01481_proC** | SP0933 |  | -6.049 |
| Spn_01482_tmk | SP0935 |  | 3.405 |
| Spn_01483_holB | SP0936 |  | 3.232 |
| Spn_01484 | SP0937 |  | 2.899 |
| Spn_01485_rsmI | SP0938 |  | 3.140 |
| Spn_01518_prsA_2 | SP0981 |  | -1.736 |
| Spn_01548_asd | SP1013 | 2.372 |  |
| Spn_01549_dapA | SP1014 | 2.242 |  |
| Spn_01560 | SP1027? |  | -1.812 |
| Spn_01567 | SP1582? | 1.603 |  |
| Spn_01721_lacR | SP1182 | -1.521 |  |
| Spn_01722 | SP1183 | -1.933 |  |
| Spn_01723_lacG | SP1184 | -2.670 | -2.042 |
| Spn_01724_lacE_2 | SP1185 | -2.466 | -2.015 |
| Spn_01725_lacF_2 | SP1186 | -1.886 | -2.033 |
| Spn_01726_lacT | SP1187 | -1.801 | -1.823 |
| Spn_01728_lacD | SP1190 | -1.516 |  |
| Spn_01753_nirC | SP1215 | -1.690 |  |
| Spn_01763_mutY | SP1228 | -2.954 |  |
| **Spn_01764_fhs** | SP1229 | -5.591 |  |
| Spn_01765_coaBC_1 | SP1230 | 2.241 |  |
| Spn_01766_coaBC_2 | SP1231 | 1.851 |  |
| Spn_01838_gdhA | SP1306 | 1.723 |  |
| Spn_01860_ABC-N/P | SP1357 | 1.650 |  |
| Spn_01861_msbA_3 | SP1358 | 1.663 |  |
| Spn_01884_amyS | SP1382 |  | 1.725 |
| Spn_01924 | SP1422 | -1.840 |  |
| Spn_01926 | SP1424 | -1.742 |  |
| Spn_02062_cshA_2 | SP1586 | 1.614 |  |
| Spn_02067_pepQ | SP1591 |  | -1.656 |
| Spn_02087 | SP1611 | 2.083 |  |
| Spn_02088 | SP1612 | 1.713 |  |
| Spn_02153_ugpA_2 | SP1682 | -1.645 |  |
| Spn_02154_yesO_1 | SP1683 | -1.695 |  |
| Spn_02155_ptsG_2 | SP1684 | -1.882 |  |
| Spn_02162_tabA_2 | SP1680 | -1.693 | -1.529 |
| Spn_02164_nanA_5 | SP1692-93 | -2.061 |  |
| Spn_02167_cah | sp1695 | -1.653 | -1.587 |
| Spn_02191_yhaP | SP1716 | -1.674 | -2.009 |
| Spn_02192_ecsA_5 | SP1717 | -1.715 | -1.983 |
| Spn_02226_sstT | SP1717 | 1.735 |  |
| Spn_02227 |  | 1.761 |  |

**Supplementary Table 4.** Genes showing differential expression (log_2_ fold change) between the mutant strains and the wild type 6B strain when cultured to mid-log growth phase in human serum. Genes in bold are deleted in the mutant strains (*proABC*, Spn_01479-81; *fhs*, Spn_01764).

| **BHN418 gene number and name** | **TIGR4 gene number**  **(if known)** | ***∆fhs* log_2_ fold change v. wild type** | ***∆proABC* log_2_ fold change v. wild type** |
| --- | --- | --- | --- |
| Spn_00015 | SP0498 | 1.683 |  |
| Spn_00022 | SP0509 |  | 1.549 |
| Spn_00023_hsdS_3 | SP0508 | 1.632 | 1.930 |
| Spn_00070 | SP1802 |  | 1.536 |
| Spn_00080 | - | 1.592 |  |
| Spn_00085_trpA | SP1811 |  | 1.552 |
| Spn_00086_trpB | SP1812 |  | 1.940 |
| Spn_00087_trpF | SP1813 |  | 1.868 |
| Spn_00088_trpC | SP1814 |  | 2.629 |
| Spn_00089_trpD2 | SP1815 |  | 2.890 |
| Spn_00090_trpG | SP1816 |  | 2.058 |
| Spn_00091_trpE | SP1817 |  | 3.467 |
| Spn_00115_xpt | SP1847 | 3.416 | 2.101 |
| Spn_00116_ygfU | SP1848 | 3.398 | 2.423 |
| Spn_00121_galT | SP1852 |  | 2.040 |
| Spn_00122_galK | SP1853 |  | 2.112 |
| Spn_00150_treC | SP1883 | -1.604 | 2.277 |
| Spn_00151_treP | SP1884 | -1.552 | 1.996 |
| Spn_00156_amiF | SP1887 | 2.153 |  |
| Spn_00157_amiE | SP1888 | 2.164 |  |
| Spn_00158_amiD | SP1889 | 1.953 |  |
| Spn_00159_amiC | SP1890 | 1.670 |  |
| Spn_00213_ply_1 | SP1923 | 3.128 | 2.507 |
| Spn_00214 | SP1924 | 3.311 | 2.647 |
| Spn_00215 | SP1925 | 2.926 | 2.245 |
| Spn_00216 | SP1926 | 2.590 | 2.077 |
| Spn_00235 |  |  | 2.613 |
| Spn_00236 | SP1956 |  | 2.415 |
| Spn_00237_ftsE_1 | SP1957 |  | 2.086 |
| Spn_00256_misCB | SP1975 |  | -1.640 |
| Spn_00260_cbf1 | SP1980 |  | 1.599 |
| Spn_00261 | SP1981 |  | 1.573 |
| Spn_00266 | SP1986 | 2.024 |  |
| Spn_00267_macB_1 | SP1987 | 2.414 | 1.531 |
| Spn_00268 | SP1988 | 2.367 |  |
| Spn_00269 | SP0109 | 1.960 |  |
| Spn_00317_licB_2 | SP2023 | 1.736 |  |
| Spn_00318_gmuA | SP2024 | 1.668 |  |
| Spn_00347 | SP2054 |  | -1.541 |
| Spn_00349_adh | SP2055 |  | 1.685 |
| Spn_00350_nagA | SP2056 | 1.585 |  |
| Spn_00356_marR_1 | SP2062 |  | -1.753 |
| Spn_00358 | SP2063 |  | -1.525 |
| Spn_00393_pstS_1 | SP2084 | -2.313 |  |
| Spn_00394_pstC_1 | SP2085 | -1.577 | 1.817 |
| Spn_00395_pstA_1 | SP2086 | -1.568 | 1.863 |
| Spn_00396_pstB3_1 | SP2087 |  | 2.336 |
| Spn_00397_phoU_1 | SP2088 |  | 2.166 |
| Spn_00415_malX | SP2108 |  | 2.440 |
| Spn_00416_malF | SP2109 | 1.968 | 2.888 |
| Spn_00417_malG | SP2110 | 1.510 | 2.173 |
| Spn_00434 | SP2125 | 1.841 |  |
| Spn_00437_tktC | SP2127 |  | 1.547 |
| Spn_00438_tktN | SP2128 |  | 1.532 |
| Spn_00439_ulaA_2 | SP2129 |  | 1.554 |
| Spn_00441_cmtB | SP2131 |  | 1.764 |
| Spn_00457_arcA | SP2148 | 1.896 | 1.988 |
| Spn_00458_arcB | SP2150 | 1.877 | 1.602 |
| Spn_00459_arcC | SP2151 | 1.587 | 1.592 |
| Spn_00460 | SP2152 | 1.535 |  |
| Spn_00469 | SP2160 |  | 1.844 |
| Spn_00470_manZ_1 | SP2161 |  | 1.655 |
| Spn_00471_manY_1 | SP2162 |  | 1.774 |
| Spn_00472_levE_1 | SP2163 |  | 1.838 |
| Spn_00473_PTS-EII_1 | SP2164 |  | 2.032 |
| Spn_00474_fucU | SP2165 |  | 2.251 |
| Spn_00475_fucA | SP2166 |  | 3.135 |
| Spn_00476_fucK | SP2167 |  | 3.457 |
| Spn_00486 | SP2178 |  | -1.613 |
| Spn_00490 | SP2182 |  | 2.199 |
| Spn_00491 | SP2182 |  | 2.264 |
| Spn_00494_glpO | SP2185 |  | 2.567 |
| Spn_00495_glpK | SP2186 |  | 2.805 |
| Spn_00520 | SP2206 |  | 1.560 |
| Spn_00551 | SP2236 | 1.620 |  |
| Spn_00562_trcF | SP0006 | 1.605 |  |
| Spn_00601_comA_1 | SP0042 | 1.874 |  |
| Spn_00602_comB | SP0043 | 2.152 |  |
| Spn_00603_purC | SP0044 | 3.518 | 1.877 |
| Spn_00604_purL | SP0045 | 3.857 | 2.879 |
| Spn_00605_purF | SP0046 | 3.442 | 2.601 |
| Spn_00606_purM | SP0047 | 3.320 | 2.503 |
| Spn_00607_purN | SP0048 | 3.039 | 2.099 |
| Spn_00608 | SP0049 | 2.049 |  |
| Spn_00609_purH | SP0050 | 2.588 |  |
| Spn_00610_purD | SP0051 | 2.265 | 2.055 |
| Spn_00611_purE | SP0053 | 2.102 | 1.896 |
| Spn_00612_purK | SP0054 | 2.640 | 2.477 |
| Spn_00617_bgaC | SP0060 |  | 6.648 |
| Spn_00618_PTS-EIIB | SP0061 |  | 6.303 |
| Spn_00619_PTS-EIIC | SP0062 |  | 5.680 |
| Spn_00620_manZ_2 | SP0063 |  | 5.571 |
| Spn_00621_PTS-EII_2 | SP0064 |  | 5.226 |
| Spn_00622_agaS | SP0265 |  | 5.001 |
| Spn_00623_mro | SP0266 |  | 3.978 |
| Spn_00638_rpsD | SP0085 | -1.815 |  |
| Spn_00640_ugpA_1 | SP0090 |  | 1.517 |
| Spn_00641_ycjP_2 | SP0091 |  | 1.620 |
| Spn_00659_artP_1 | SP0112 | 2.162 |  |
| Spn_00660_argG | SP0113 | 1.743 |  |
| Spn_00662_lagD_1 | SP0312 |  | 1.510 |
| Spn_00663 | SP0115 |  | 2.447 |
| Spn_00665 | SP0115 |  | 1.600 |
| Spn_00667 |  |  | 1.585 |
| Spn_00669 | SP0115 |  | 1.550 |
| Spn_00676_mnmA | SP0118 | -1.669 |  |
| Spn_00698_mutR | SP0141 | 1.550 |  |
| Spn_00699 | SP0142 | 1.536 |  |
| Spn_00700 | SP0143 | 1.595 |  |
| Spn_00701 | SP0144 | 1.873 |  |
| Spn_00702 | SP0145 | 2.426 |  |
| Spn_00712_ypdA | SP0155 |  | 1.938 |
| Spn_00716 | SP0159 | 1.797 |  |
| Spn_00728_ribA | SP0176 | -1.842 |  |
| Spn_00729_ribE | SP0175 | -2.013 |  |
| Spn_00730_ribD | SP0178 | -1.715 |  |
| Spn_00757 | SP0191 |  |  |
| Spn_00770_rpsJ | SP0208 | -1.510 | -1.574 |
| Spn_00816_gmuB | SP0249 |  | 1.915 |
| Spn_00817_gmuC_4 | SP0250 |  | 2.724 |
| Spn_00818_hpdB | SP0251 |  | 2.554 |
| Spn_00819_fsaA | SP0252 |  | 2.110 |
| Spn_00820_gldA | SP0253 |  | 1.898 |
| Spn_00832_glmS | SP0266 | -1.854 |  |
| Spn_00834_pulA_1 | SP0268 |  | 1.943 |
| Spn_00851_pbuO | SP0287 | 2.173 |  |
| Spn_00852 | SP0288 | 2.580 |  |
| Spn_00873 | SP0314 |  | 2.259 |
| Spn_00876 | SP0318 |  | 1.942 |
| Spn_00879_manX | SP0321 |  | 2.104 |
| Spn_00880_ugl | SP0322 |  | 2.576 |
| Spn_00881_levE_2 | SP0323 |  | 2.121 |
| Spn_00882_agaC | SP0324 |  | 2.016 |
| Spn_00883_manZ_4 | SP0325 |  | 2.035 |
| Spn_00915 | SP0368 |  | 2.517 |
| Spn_00964_marR_2 | SP0416 |  | 1.711 |
| Spn_00965_fabH | SP0417 |  | 1.711 |
| Spn_00966_acpP_2 | SO0418 |  | 1.570 |
| Spn_00967 | SP0419 | 3.283 | 3.043 |
| Spn_00968_fabD | SP0420 | 3.180 | 3.142 |
| Spn_00969_fabG | SP0421 | 3.206 | 3.142 |
| Spn_00970_fabF | SP0422 | 2.728 | 2.781 |
| Spn_00971_accB | SP0423 | 2.863 | 3.022 |
| Spn_00972_fabZ | SP0424 | 3.002 | 3.107 |
| Spn_00973_accC | SP0425 | 3.067 | 3.195 |
| Spn_00974_accD | SP0426 | 2.890 | 3.019 |
| Spn_00975_accA | SP0427 | 2.931 | 3.066 |
| Spn_00992_ilvB | SP0445 | 2.347 |  |
| Spn_00993_ilvH | SP0446 | 2.698 |  |
| Spn_00994_ilvC | SP0447 | 2.896 |  |
| Spn_00995 | SP0450 | 2.675 |  |
| Spn_00996 | SP0449 | 2.698 |  |
| Spn_01030_cbiQ | SP0483 | 1.780 | 1.896 |
| Spn_01078_spiR2_1 | SP0526 | -1.758 |  |
| Spn_01082_dnaK_3 | SP0517 |  | -4.042 |
| Spn_01089 | SP0524 |  | 2.786 |
| Spn_01094_lcnD_1 | SP0529 |  | 2.024 |
| Spn_01096_lagD_2 |  |  | 2.202 |
| Spn_01097_blpA2 |  | 2.380 | 3.702 |
| Spn_01098_blpI | SP0531 |  | 3.391 |
| Spn_01099 | SP0532 | 2.814 | 4.556 |
| Spn_01100_blpN_1 | SP0533 | 2.800 | 4.650 |
| Spn_01102 | SP0536 | 1.763 | 3.312 |
| Spn_01108_blpX | SP0544 | 1.917 | 3.826 |
| Spn_01109_pncO | SP0545 | 3.024 | 4.701 |
| Spn_01110_blpZ | SP0546 | 2.397 | 4.284 |
| Spn_01111 | SP0547 | 2.438 | 3.871 |
| Spn_01114_rimP | SP0552 |  | -1.605 |
| Spn_01162_yecS_2 | SP0607 |  | 1.522 |
| Spn_01192_prtP | SP0641 | 2.073 | 2.460 |
| Spn_01193 | SP0645 |  | 5.451 |
| Spn_01194 | SP0646 |  | 5.173 |
| Spn_01195_gatC_2 | SP0647 |  | 5.105 |
| Spn_01196 |  |  | 4.106 |
| Spn_01197_lacZ | SP0648 |  | 4.517 |
| Spn_01230_typA | SP0681 |  | -1.733 |
| Spn_01232 | SP0684/685 |  | 2.998 |
| Spn_01233 | SP0686 |  | 1.778 |
| Spn_01234 | SP0686 |  | 2.791 |
| Spn_01235_glnQ_2 | SP0709 |  | 2.392 |
| Spn_01240_pyrF | SP0701 | -1.735 |  |
| Spn_01241_pyrE | SP0702 | -1.728 |  |
| Spn_01242 | SP0703 |  | 2.067 |
| Spn_01243 | SP0704 |  | 2.262 |
| Spn_01244 | SP0705 |  | 1.669 |
| Spn_01245 | SP0706 |  | 1.830 |
| Spn_01246_cysA | SP0707 |  | 1.895 |
| Spn_01254_lctO | SP0715 |  | 1.546 |
| Spn_01302_livH | SP0750 | 1.803 |  |
| Spn_01303_livM | SP0751 | 1.992 |  |
| Spn_01304_lptB | SP0752 | 2.277 |  |
| Spn_01305_livF | SP0753 | 2.262 |  |
| Spn_01306 | SP0754 | 1.845 |  |
| Spn_01324_rpsP_2 | SP0775 | -1.751 |  |
| Spn_01325 | SP0776 | -1.546 |  |
| Spn_01332_bioY2 | SP0783 | 3.851 | 2.194 |
| Spn_01381_coaA | SP0839 |  | 1.718 |
| Spn_01384_deoA | SP0842 |  | 1.604 |
| Spn_01400_sarA_1 | SP0857 | 1.526 |  |
| Spn_01401 | SP0858 |  |  |
| Spn_01402 | SP0859 | 1.721 |  |
| Spn_01403_pcp_2 | SP0860 | 1.925 |  |
| Spn_01427 | SP0879 |  | -1.780 |
| Spn_01456 | SP0910 | -2.398 | -2.029 |
| Spn_01457 | SP0911 |  | 1.582 |
| Spn_01479_proB | SP0931 |  | -5.550 |
| Spn_01480_proA | SP0932 |  | -5.456 |
| Spn_01481_proC | SP0933 |  | -5.189 |
| Spn_01482_tmk | SP0935 |  | 3.649 |
| Spn_01483_holB | SP0936 |  | 4.023 |
| Spn_01484 | SP0937 |  | 3.808 |
| Spn_01485_rsmI | SP0938 |  | 4.087 |
| Spn_01495_kpsT | SP0957 |  | 3.604 |
| Spn_01496 |  |  | 3.964 |
| Spn_01501_pyrDII | SP0963 | -2.361 |  |
| Spn_01502_pyrD | SP0964 | -1.638 |  |
| Spn_01514_tehB | SP0977 | 1.543 |  |
| Spn_01518_prsA_2 | SP0985 |  | -1.810 |
| Spn_01542_hemH | SP1009 |  | 1.770 |
| Spn_01548_asd | SP1013 | 2.089 |  |
| Spn_01549_dapA | SP1014 | 1.937 |  |
| Spn_01632_gapN | SP1119 |  | -2.254 |
| Spn_01633_glgB | SP1121 |  | 3.092 |
| Spn_01634_glgC | SP1122 | 1.851 | 2.915 |
| Spn_01635_glgD | SP1123 | 1.864 | 2.755 |
| Spn_01636_glgA | SP1124 | 1.692 | 2.447 |
| Spn_01672 |  | -2.602 | -2.166 |
| Spn_01673 |  |  | -1.511 |
| Spn_01674 |  |  | -1.686 |
| Spn_01677_macB_6 | SP0912 | -1.533 | -1.847 |
| Spn_01678 | SP0913 |  | -1.526 |
| Spn_01718_nrdH | SP1178 |  | -1.780 |
| Spn_01727 | SP1189 |  | 2.391 |
| Spn_01728_lacD | SP1190 |  | 2.987 |
| Spn_01729_lacC-2 | SP1191 |  | 3.101 |
| Spn_01730_lacB-2 | SP1192 |  | 3.097 |
| Spn_01731_lacA | SP1193 |  | 3.213 |
| Spn_01736 | sp1198 |  | 4.378 |
| Spn_01753_nirC | SP1215 |  | 1.582 |
| Spn_01763_mutY | SP1228 | -4.655 |  |
| Spn_01764_fhs | SP1229 | -5.698 |  |
| Spn_01765_coaBC_1 | SP1230 | 4.255 |  |
| Spn_01766_coaBC_2 | SP1231 | 4.067 |  |
| Spn_01767_panT | SP1699 | 4.049 |  |
| Spn_01780_ybjI | SP1247 |  | 1.772 |
| Spn_01781_smc | SP1248 |  | 1.765 |
| Spn_01784_guaC | SP1249 |  | 1.705 |
| Spn_01799_licC | SP1267 | 2.442 |  |
| Spn_01800 | SP1268 | 2.391 |  |
| Spn_01801 | SP1269 | 2.537 |  |
| Spn_01802_idnD | SP1270 | 2.269 |  |
| Spn_01803_tarI | SP1271 | 2.256 |  |
| Spn_01819_pyrP | SP1286 | -2.305 |  |
| Spn_01840_hlyB_2 |  | 1.654 |  |
| Spn_01841 |  | 1.643 |  |
| Spn_01884_amyS | SP1382 |  |  |
| Spn_01896_glnH | SP1394 |  | -2.106 |
| Spn_01917_nagB | SP1415 | 1.964 |  |
| Spn_01919_lytA_5 | SP1416 | 2.298 |  |
| Spn_01920 | too short | 1.787 |  |
| Spn_01921_ydaF_4 | SP1419 | 1.914 |  |
| Spn_01955_pdx1 | SP1468 |  | -1.617 |
| Spn_01957_apbE | SP1470 |  | 1.524 |
| Spn_01958_azr_1 | SP1471 |  | 1.533 |
| Spn_02005_lmrA_2 | too short | 3.335 |  |
| Spn_02006_sarA_2 | SP1527 | 2.835 |  |
| Spn_02012_murE2 | SP1530 | -1.749 |  |
| Spn_02057_rebM | SP1578 |  | 1.842 |
| Spn_02058_ugpC | SP1580 |  | 1.986 |
| Spn_02060_yecD_1 | SP1583 |  | 1.733 |
| Spn_02063_yhjX | SP1587 | 2.337 |  |
| Spn_02086 | SP1611 | 3.162 |  |
| Spn_02087 | SP1612 | 2.123 |  |
| Spn_02088 | SP1612 | 2.827 |  |
| Spn_02101_rpsO_2 | SP1626 |  | -1.736 |
| Spn_02120_psaA | SP1650 | 1.673 |  |
| Spn_02128 | SP1658 |  | -1.775 |
| Spn_02146_bglK_2 | SP1675 | 1.589 | 2.780 |
| Spn_02148 | SP1677 | 2.395 | 2.777 |
| Spn_02149 | SP1678 | 2.332 | 3.005 |
| Spn_02150 | SP1679 | 2.358 | 3.350 |
| Spn_02151_tabA_1 | SP1680 | 2.418 |  |
| Spn_02152_ycjP_4 |  | 2.042 | 3.042 |
| Spn_02153_ugpA_2 | SP1682 | 2.288 | 3.678 |
| Spn_02154_yesO_1 | SP1683 | 1.825 | 3.496 |
| Spn_02155_ptsG_2 | SP1684 | 1.679 | 4.381 |
| Spn_02156_nanE2 | SP1685 | 1.546 | 4.044 |
| Spn_02164_nanA_5 | SP1692-93 |  | 2.910 |
| Spn_02165 | SP1694 |  | 2.316 |
| Spn_02167_cah | SP1695 | -2.703 | -2.359 |
| Spn_02181 | SP1707 |  | 2.502 |
| Spn_02182 | SP1708 |  | 2.488 |
| Spn_02227 | SP1754 | 2.448 |  |
| Spn_02228 |  | 2.109 |  |

**Supplementary Table 5. Mean values and standard deviation for 18 *in vitro* conditions and two *in vivo* niches obtained from van Opijnen and Camilli (4).** Values with an asterisk are statistically significant.

| ***Mutant phenotype data*** | | | | |
| --- | --- | --- | --- | --- |
| **Growth condition** | ***SP1229*** | ***SP0931*** | ***SP0932*** | ***SP0933*** |
| Galactose  Fructose  GlnNac  Cellobiose  Raffinose  Sucrose  Glucose  Maltose  Mannose  MMS  pH6  Temperature  H2O2  MMS  Norfloxacin  Lung  Sialic acid  Transformation  Bipyridyl  Nasopharynx | 0.87(0.06)*  0.89(0.03)*  0.90(0.05)*  0.77(0.09)*  0.88(0.06)*  0.84(0.08)*  0.87(0.06)*  0.84(0.08)*  1.13(0.06)*  0.85(0.07)  0.77(0.09)*  0.85(0.07)*  0.91(0.09)*  0.85(0.07)*  0.90(0.04)*  0.42(0.45)*  0.98(0.06)  1.02(0.06)  0.97(0.07)  1.08(0.09) | 0.98(0.04)  0.98(0.03)  0.98(0.03)  1.02(0.05)  1.00(0.04)  0.93(0.09)  0.98(0.04)  0.97(0.05)  1.02(0.04)  1.00(0.05)  1.01(0.06)  0.97(0.06)  0.98(0.05)  1.00(0.05)  0.96(0.05)  0.66(0.38)*  1.03(0.07)  1.06(0.09)  0.98(0.06)  1(0.10) | 1.00(0.07)  0.98(0.03)  0.99(0.03)  1.02(0.05)  1.02(0.08)  0.94(0.06)  1.00(0.07)  0.99(0.05)  1.04(0.09)  0.98(0.07)  1.00(0.04)  0.98(0.05)  0.98(0.07)  0.98(0.07)  1.01(0.10)  0.51(0.45)*  1.02(0.06)  1.06(0.11)  0.99(0.07)  1.10(0.07) | 1.00(0.04)  0.89(0.02)  0.98(0.03)  1.01(0.05)  1.01(0.06)  0.97(0.05)  1.00(0.04)  1.00(0.03)  1.03(0.05)  1.02(0.05)  1.04(0.02)  1.01(0.04)  0.97(0.04)  1.02(0.05)  1.01(0.05)  0.42(0.44)*  1.04(0.06)  1.07(0.06)  1.01(0.06)  1.13(0.05) |

**References:**

1.	Martin B, Prudhomme M, Alloing G, Granadel C, Claverys JP. 2000. Cross-regulation of competence pheromone production and export in the early control of transformation in Streptococcus pneumoniae. Mol Microbiol 38:867-78.

2.	Liu X, Gallay C, Kjos M, Domenech A, Slager J, van Kessel SP, Knoops K, Sorg RA, Zhang JR, Veening JW. 2017. High-throughput CRISPRi phenotyping identifies new essential genes in Streptococcus pneumoniae. Mol Syst Biol 13:931.

3.	Granok AB, Parsonage D, Ross RP, Caparon MG. 2000. The RofA binding site in Streptococcus pyogenes is utilized in multiple transcriptional pathways. J Bacteriol 182:1529-40.

4.	van Opijnen T, Camilli A. 2012. A fine scale phenotype-genotype virulence map of a bacterial pathogen. Genome Res 22:2541-51.
